# Physiological synaptic activity and recognition memory are fueled by astroglial glutamine

**DOI:** 10.1101/2020.10.06.328823

**Authors:** Giselle Cheung, Danijela Bataveljic, Naresh Kumar, Julien Moulard, Glenn Dallérac, Josien Visser, Astrid Rollenhagen, Cédric Mongin, Oana Chever, Alexis-Pierre Bemelmans, Joachim Lübke, Isabelle Leray, Nathalie Rouach

## Abstract

Presynaptic glutamate replenishment is fundamental to brain function. In high activity regimes, such as epileptic episodes, this process is thought to rely on the glutamate-glutamine cycle between neurons and astrocytes. However the presence of an astroglial glutamine supply, as well as its functional relevance *in vivo* in the healthy brain remain controversial, partly due to a lack of tools that can directly examine glutamine transfer. Here, we generated a novel fluorescent probe that tracks glutamine in live cells, which provided direct visual evidence of an activity-dependent glutamine supply from astroglial networks to presynaptic structures under physiological conditions. This mobilization is mediated by connexin43, an astroglial protein with both gap-junction and hemichannel functions, and is essential for synaptic transmission and object recognition memory. Our findings uncover an indispensable recruitment of astroglial glutamine in physiological synaptic activity and memory via an unconventional pathway, thus providing an astrocyte basis for cognitive processes.

## Introduction

Neurotransmitter replenishment upon release of synaptic vesicles is essential for many neuronal functions. Although some neurotransmitters can be retrieved by neurons through various reuptake mechanisms, glutamate is mainly removed by perisynaptic astrocytes via glutamate transporters^1,2^. This removal of glutamate not only helps prevent excitotoxicity, but also acts as a local mechanism to recycle synaptic glutamate. The glutamate-glutamine cycle postulates that astrocytes take up glutamate from the synaptic cleft and transform it into glutamine, which is transported back to neurons to be converted to glutamate^3^. Such a mechanism would be important *in vivo* because *de novo* glutamine synthesis does not occur at nerve terminals^4,5^. In fact, it was estimated that, without replenishment, the pool of presynaptic glutamate would be exhausted within a minute of basal synaptic activity^6^. While this cycle has been implicated in epilepsy, a hyperactive state where the rate of glutamate release is greatly enhanced^7–9^, its relevance to physiological conditions is still under debate due to conflicting reports. Early reports claimed that inhibition of glutamate to glutamine conversion or glutamine transport into neurons does not affect excitatory synaptic transmission or quantal size^10–12^. Chronic inhibition of glutamine synthetase was also found to have no effect on learning and memory in mice^13^. In contrast, other studies showed that presynaptic glutamine transport contributes to miniature EPSC amplitude^14^ and astroglial glutamine synthesis sustains excitatory transmission^15^. A major step forward in helping to clarify the role of astrocytic glutamine would be to develop tools capable of directly assessing glutamine transfer. While radiolabeled-glutamine has previously been used for quantitative measurements of glutamine in brain samples^16,17^, this approach offers poor spatial and temporal resolution. More recently, two genetically encoded FRET-sensors for glutamine have been developed for plant root tips^18^ and cos7 cells^19^. Although useful in live preparations, these sensors depend on genetic manipulations, and cannot distinguish a directional flow of glutamine across adjacent cells. In the absence of tools that can track directional glutamine movement, a direct role for astroglial glutamine supply during physiological synaptic activity remains unclear. Here we develop a novel glutamine fluorescent probe, which allowed us to gather direct visual evidence for activity-dependent astrocyte-neuron glutamine transfer under physiological conditions. Further, we find that the role of glutamine is dependent on astroglial connexin 43 hemichannels, and provide evidence for the functional relevance of astroglial glutamine in physiological synaptic transmission and cognition.

## Results

### Generation of a fluorescent glutamine molecule

We generated a novel fluorescent rhodamine-tagged glutamine molecule (RhGln) as a probe to directly visualize glutamine in live cells from brain tissues. RhGln was synthesized in a 5-step procedure (Fig. 1a and Supplementary Fig. 1), and is a small (0.64kDa), brightly fluorescent, stable, water soluble and non-hydrolysable glutamine analog. The fluorescent tag was conjugated to the amide side chain to prevent hydrolysis of glutamine to glutamate by glutaminase, as previously described for theanine^20^, thereby enabling visualization of glutamine only but not its metabolites. Further characterization showed that RhGln has an absorption and emission maxima at 580 and 601nm, respectively (Fig. 1b-c). Compared to the unconjugated rhodamine molecule (Rh101), RhGln showed a slight bathochromic shift of absorption and emission spectra and a fluorescence emission quantum yield of 0.6 (Fig. 1d-e).

**Fig. 1.**
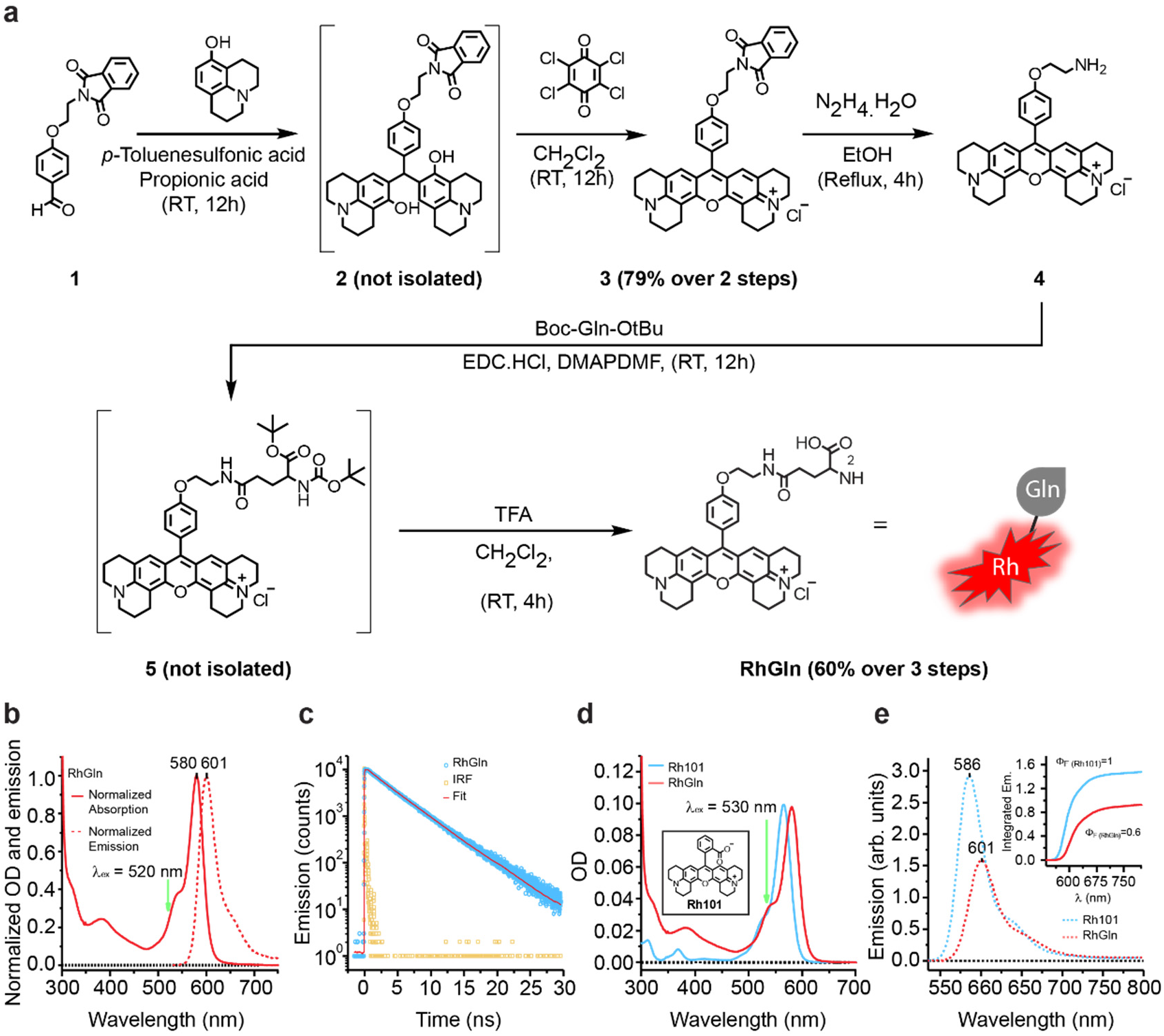
Synthesis and characterization of fluorescent rhodamine-tagged glutamine (RhGln) molecule. **a,** Fluorescent RhGln molecule was prepared using a 5-step chemical synthesis with 48% overall yield. **b,** Characterization of steady state electronic absorption (solid line) and emission (dotted line) spectra of RhGln at excitation wavelength (λ_ex_) of 520nm in intracellular solution at 20°C, showing sharp absorption and emission maxima of 580 and 601 nm, respectively. **c,** Fluorescence lifetime decay of RhGln in intracellular solution at λ_ex_ of 520nm (blue circles). The instrument response function (IRF, orange squares) and fitted line (Fit, red) are also shown. **d, e,** Comparison of absorption **(d)** and emission **(e)** spectra between RhGln (red line, 0.83μM) and its rhodamine precursor (Rh101, blue line, 0.94μM) at an λ_ex_ of 530nm in intracellular solution at 20°C is shown with a relative fluorescence emission quantum yield value (Φ_F_) of 0.6 for RhGln (inset).

### Activity-dependent mobilization of glutamine from astrocytes to presynaptic structures

To visualize cellular transfer of astroglial glutamine, we dialyzed RhGln (0.8mM, 20min) into single CA1 astrocytes from hippocampal slices using the whole cell patch-clamp technique, and observed diffusion of RhGln into neighboring cells (Fig. 2a), identified as GFP-positive astrocytes from Aldh1l1-eGFP mice (Fig. 2b). RhGln labeling in neighboring astrocytes was located in soma and processes, including both main and fine processes, as revealed by RhGln dialysis of GFAP-eGFP astrocytes (Fig. 2c). RhGln-labeling was not due to extracellular uptake, as RhGln was delivered selectively and intracellularly to single astrocytes via a patch pipette and extracellular bulk loading of slices with RhGln did not label astrocytes (Supplementary Fig. 2). The diffusion of RhGln into astroglial networks occurred via gap junction (GJ) channels, because it was inhibited by the GJ blocker carbenoxolone (CBX, Fig. 2a and 2e). We next investigated whether the transfer of astroglial glutamine was activity-dependent by comparing RhGln distribution during basal and evoked activity triggered by Schaffer collaterals repetitive stimulation (10Hz, 30s). The stimulation evoked field excitatory postsynaptic potentials (fEPSPs) and an astrocytic response (Fig. 2d). Surprisingly, we found that glutamine intercellular transfer was reduced in astrocytes upon stimulation (Fig. 2a and 2e), implying that synaptic activity induces a redistribution of glutamine away from astroglial networks. We examined where glutamine was redistributed to at a subcellular level, and found that RhGln did not only spread into neighboring astrocytes, but was also present in punctate structures surrounding the patched astrocyte (Fig. 2f). Furthermore, the RhGln punctate labeling was much stronger and spread further upon stimulation (Fig. 2f, 2g and 2j), indicating an activity-dependent mobilization of glutamine away from the astrocyte soma. This activity-dependent punctate labeling was specific to RhGln as it was not observed with the rhodamine (Rh) dye alone (Fig. 2f, 2h and 2j). It is also unlikely to result from activity-dependent morphological changes in astrocytes, since we found no change in soma size, domain area and ramifications, as determined in GFP- and GFAP-positive astrocytes from Aldh1l1-eGFP mice (Supplementary Fig. 3). Because fine astroglial processes are often found near synaptic structures^21^ and activity-dependent RhGln punctate labeling was inhibited by blocking neuronal glutamine transport with methylaminoisobutyric acid (MeAIB, Fig. 2f, 2i and 2j), we investigated whether RhGln entered synaptic compartments upon activity. Confocal microscopy indicated that the extended punctate labeling showed increased co-localization with VGlut1, a glutamatergic presynaptic marker (Fig. 3a-b). To unequivocally identify the cellular structure of the RhGln punctate labeling, we used super-resolution STED imaging of RhGln-filled hippocampal GFAP-eGFP-positive astrocytes and confirmed that these punctate structures were indeed presynaptic (Fig. 3c-e). Together, our findings identify an activity-dependent mobilization of astroglial glutamine from astroglial networks into adjacent presynaptic structures.

**Fig. 2.**
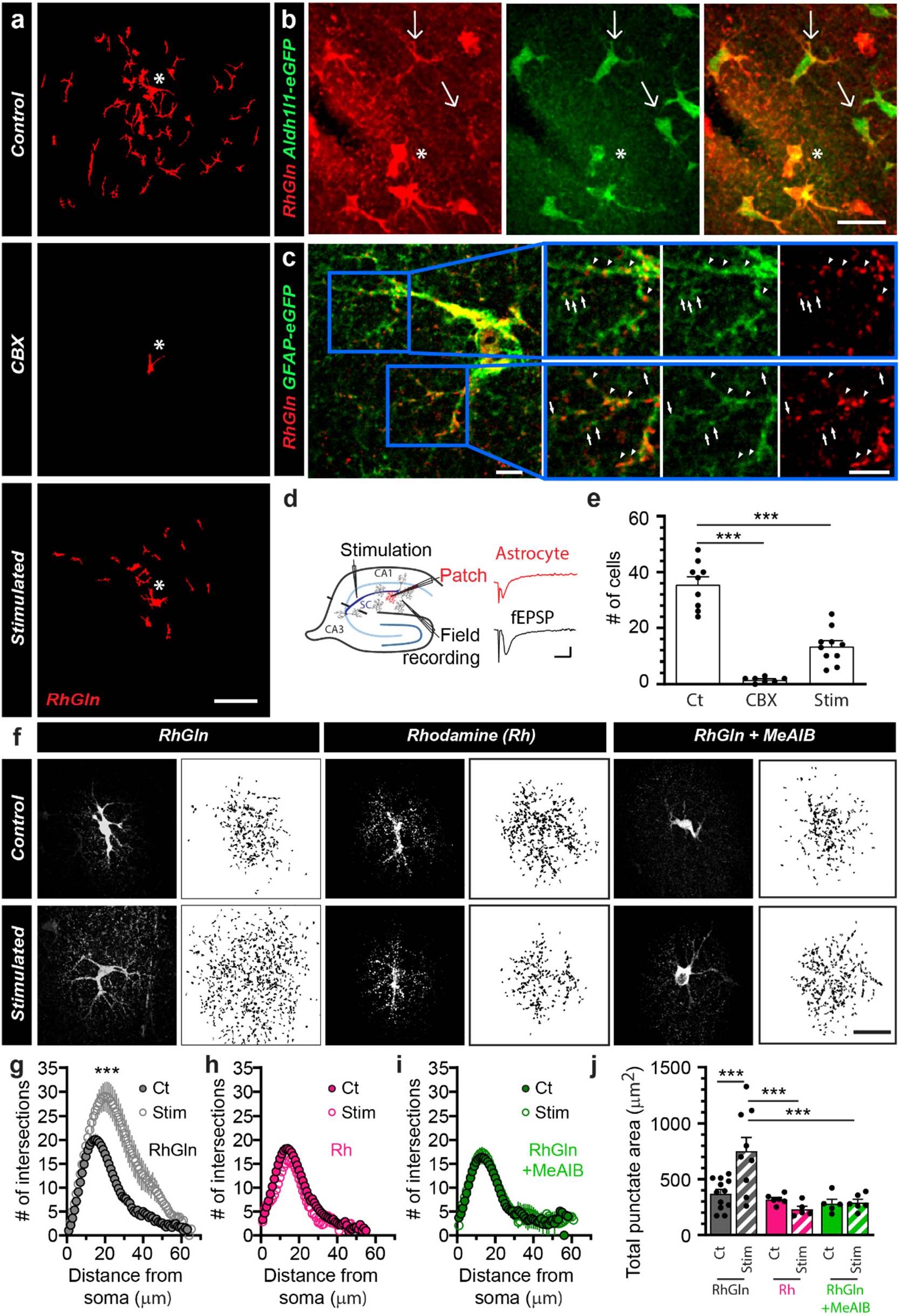
RhGln reveals activity-dependent redistribution of glutamine away from astroglial networks to subcellular punctate structures. **a,** RhGln traffics through gap junction-mediated astroglial networks when dialyzed (0.8mM, 20min) into a single CA1 hippocampal astrocyte via the patch pipette. The representative images illustrate the transfer of RhGln from the patched astrocyte (asterisks) to neighboring cells under control condition, which was abolished by the gap-junction blocker carbenoxolone (CBX, 200μM), and reduced by repetitive synaptic stimulation (10Hz, 30s every 3min for 20min). **b,** RhGln under control conditions was found distributed in neighboring cells identified as Aldh1l1-positive astrocytes, indicating the spread of glutamine into astroglial networks. Asterisks mark the positions of the patched cells. Thin arrows mark astrocytes in which RhGln labeling is found primarily in processes. **c,** RhGln puncta were observed along main (arrow heads) and fine processes (arrows) of a GFAP-positive cell. Higher magnification of two areas (blue boxes) are also shown. **d,** Schematic and sample traces showing simultaneous recordings of evoked depolarization of a patched astrocyte (Patch, red trace) and field excitatory postsynaptic potential (fEPSP, Field recording, black trace) during stimulation of the Schaffer collaterals (Stimulation). **(e)** Quantification of RhGln transfer from the patched astrocyte to neighboring cells under the different conditions (Control, Ct, n=9; CBX, n=7, *p*<0.0001; Stimulated, Stim, n=10, *p*<0.0001, One-way ANOVA with Bonferroni’s post hoc test). **f**, Punctate RhGln-labeling surrounding dye-filled astrocytes is enhanced by stimulation as shown in representative confocal (dark background) and thresholded binary (white background) images **(f left)**, quantification by Sholl analysis (Ct, n=12; Stim, n=9, *p*<0.0001, Two-way ANOVA in **g)** and total punctate area (*p*=0.0003 between Ct and Stim, One-way ANOVA with Bonferroni’s post hoc test in **j**). This was not observed when the unconjugated rhodamine dye was used (Rh, 0.8mM in **f middle**: Ct, n=6; Stim, n=5, *p*=0.7012 in **h**; and *p*>0.9999 between Ct and Stim with Rh, and *p*<0.0001 between Stim of RhGln and Rh in **j)** or in the presence of MeAIB, an antagonist of neuronal glutamine uptake (RhGln + MeAIB, 20mM in **f right**: Ct, n=5; Stim, n=6, *p*=0.7999 in **i**; p>0.9999 between Ct and Stim with RhGln + MeAIB, and *p*=0.0002 between Stim of RhGln and RhGln + MeAIB in **j**); Two-way ANOVA was used in **b** and One-way ANOVA with Bonferroni’s post hoc test in **c.** Scale bars: **a,** 50μm; **b and f,** 20μm; **c,** 5μm; **d,** 0.5mV (Astrocyte), 0.2mV (fEPSP), 20ms. Asterisks indicate statistical significance (***: *p*<0.0001).

**Fig. 3.**
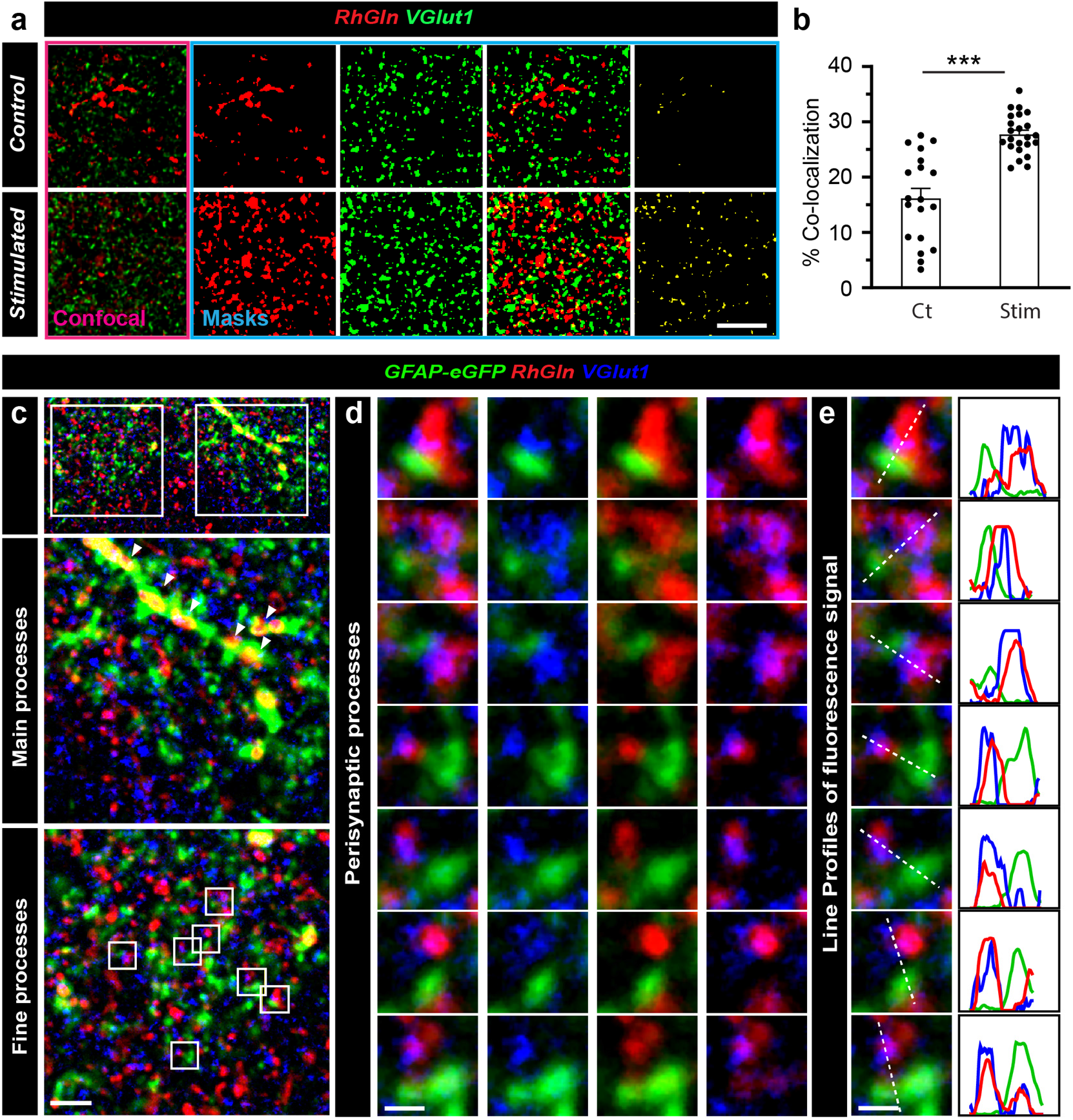
Astroglial RhGln enters presynaptic compartments upon synaptic activity. **a,** Sample confocal (pink box) and binary images (blue box) showing that after dialysis of a single astrocyte with RhGln (0.8mM, 20min), RhGln-labeled puncta (red) are observed and show increased co-localization with the glutamatergic presynaptic marker VGlut1 (green) under repetitive synaptic stimulation (10Hz, 30s every 3min for 20min). **b,** Bar graph (Mean ± SEM) showing % co-localization normalized to total area of RhGln-filled structures for Ct (n=19 fields, 3 independent experiments) and Stim (n=23 fields, 3 independent experiments, *p*<0.0001, unpaired Student’s t-test) conditions. **c-d,** STED super-resolution imaging confirmed co-localization of VGlut1 (blue) with RhGln (red) found directly next to an astroglial compartment (green) after dialysis of a GFAP-eGFP expressing astrocyte (green) with RhGln (0.8mM, 20min) during repetitive synaptic stimulation, as shown in the sample images **(c)**. Higher magnifications of regions containing either main (middle panel) or fine (lower panel) astroglial processes are shown. Arrowheads denote accumulation of RhGln along a main process. **d,** Representative images containing perisynaptic astroglial processes marked by boxes in **c** (lower panel). **e,** Line profiles measured along individual white dotted lines for individual channels to illustrate co-localization. Scale bars: **a,** 5μm; **c,** 2μm; **d** and **e**, 0.5μm. Asterisks indicate statistical significance (***: *p*<0.0001).

### Astroglial connexin 43 is expressed in perisynaptic astroglial processes and displays activity-dependent hemichannel opening

To track down the mechanism involved in activity-dependent glutamine supply by astrocytes, we focused on astroglial proteins that have regulatory roles in both the astroglial network and at a perisynaptic level. Astroglial connexins (Cxs) have both GJ and hemichannel (HC) functions, mediating the transfer of small molecules within and out of the astroglial network, respectively^22^. This unique property may drive the redistribution of glutamine from the astroglial network into perisynaptic compartments and then synapses. While astrocytes abundantly express both Cx30 and 43 isoforms, evidence suggests that only Cx43 regulates physiological synaptic activity via its HC function^22,23^. However, to establish the involvement of Cx43 in glutamine supply, two prerequisites have to be fulfilled: expression in close proximity to synapses, and activity-dependent regulation of HC activity. To assess Cx43 localization, we used immunolabeling and found that Cx43 protein is expressed throughout hippocampal astrocytes including fine processes away from the cell soma and close to VGlut1-positive glutamatergic presynapses (Fig. 4a). To further support this localization, we co-purified perisynaptic astroglial processes with hippocampal synaptosomes^24,25^ and found an enrichment of Cx43 proteins in synaptosomal fractions compared to total hippocampal extracts (Fig. 4b-c). Preparations obtained from glial conditional Cx43 knockout mice (-/-) lacked Cx43 protein expression, confirming that this protein enrichment is of glial origin. Lastly, at the ultrastructural electron microscopic level, we observed immunogold particles labeling Cx43 in small astroglial processes close to synaptic complexes, with a minimum distance of 72.2 nm and an average distance of 265.8 ± 115.5 nm to the nearest active zone (Fig. 4d-e). To test whether physiological synaptic stimulation alters Cx43 HC activity, we performed ethidium bromide (EtBr) uptake assays in acute slices (Fig. 4f), and observed that synaptic stimulation (10Hz, 30s) markedly increased EtBr uptake. This uptake was mediated by Cx43 HCs, as it was abolished by the Gap26 peptide, a specific blocker of Cx43 HCs, but not by a scrambled peptide (Gap26Sr, Fig. 4f-h). Altogether, these data reveal that astroglial Cx43 is located near synapses and that its HC function is activity-dependent.

**Fig. 4.**
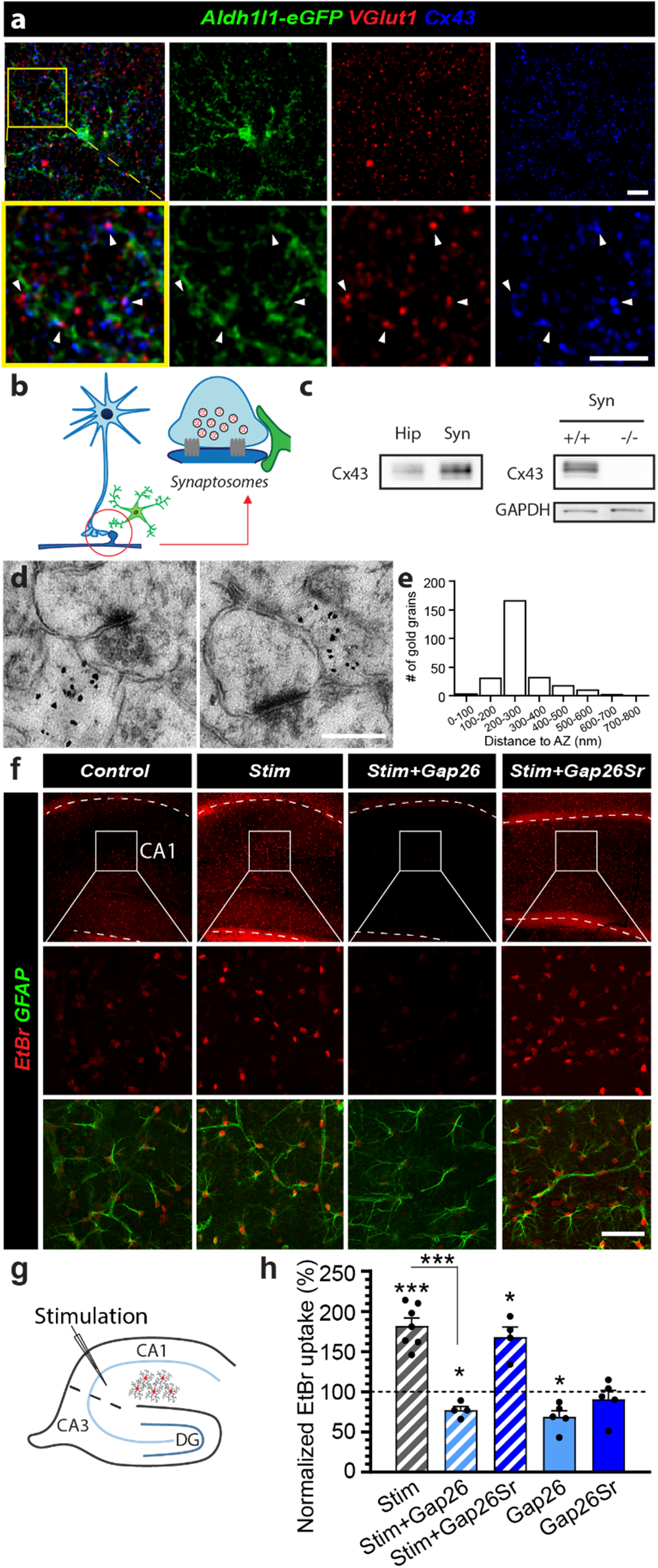
Cx43 is expressed in perisynaptic astroglial processes and displays enhanced hemichannel function upon physiological synaptic activity. **a,** Representative confocal images showing close proximity of Cx43 (blue) in Aldh1l1-eGFP positive astrocytes (green) to presynaptic structures immunolabeled for VGlut1 (red). Higher magnification images of a region containing astroglial processes (yellow square) are shown in the bottom row. Arrowheads denote points of close contact. **b,** Schematic illustration of co-purification of perisynaptic astroglial processes in crude synaptosomes. **c,** Representative western blots showing an enrichment of Cx43 protein in synaptosomal preparations (Syn) compared to total hippocampal lysates (Hip) in wild type (+/+), but not in glial conditional Cx43 knockout (-/-) mice. GAPDH was used as a loading control. **d,** Representative high magnification electron micrographs showing the presence of Cx43 protein labeled by immunogold particles in astroglial processes near synaptic complexes. **e,** Distribution histogram of distance between Cx43 gold grains and the nearest active zone. **f,** Sample images of ethidium bromide (EtBr, red) uptake in hippocampal GFAP-immunolabeled astrocytes (green) are shown in different conditions: Control, Stim (10Hz, 30s every 3min for 20min) in the absence or presence of the Cx43 HC blocker Gap26 or a Gap26 scramble version (Gap26Sr). Higher magnifications of the CA1 stratum radiatum subregion are shown in bottom two rows. **g**, Schematic illustrating stimulation of hippocampal Schaffer collaterals and EtBr uptake in neighboring astrocytes. **h**, Quantification of EtBr uptake normalized to 100% control (dotted line) is shown. Stimulation enhanced EtBr uptake by nearly 2-fold (Control, n=6; Stim, n=7, *p*=0.0002 between Control and Stim, One-sampled t-test). This enhanced uptake was not observed in the presence of Gap26 (Stim+Gap26, n=4, *p*<0.0001 between Stim and Stim+Gap26, *p*=0.0168 with control) but persisted with Gap26Sr (Stim+Gap26Sr, n=4, *p*=0.5681 between Stim and Stim+Gap26Sr, *p*=0.0125 with control), while Gap26 alone decreases EtBr uptake from control level (n=5, *p*=0.014) but not Gap26Sr (n=5, *p*=0.4443). One-sampled t-test was used to compare each bar to 100% control and One-way ANOVA with Sidak’s post hoc test was used to compare between bars. Scale bars: **a,** 5μm, **d,** 0.5μm (left), 0.3μm (right); **f,** 450μm (top), 50μm (middle and bottom). Asterisks indicate statistical significance (***: *p*<0.001; *: *p*<0.05).

### Connexin 43 mediates activity-dependent transfer of glutamine from hippocampal astrocytes to synapses

Next, we tested whether Cx43 HCs mediate glutamine supply from astrocytes to synapses in an activity-dependent manner. To do so, we either genetically disrupted Cx43 using glial conditional Cx43 knockout mice (-/-), or acutely inhibited Cx43 HC activity with Gap26 treatment. In astrocytes lacking Cx43 (-/-), the diffusion of RhGln was not only decreased under basal conditions, but also became insensitive to synaptic stimulation (Fig. 5a-b). The same effects were observed with the Gap26 but not with the Gap26Sr peptide, indicating that activity-dependent glutamine transfer from astrocytes to synapses is mediated by Cx43 HCs (Fig. 5a-b). Quantitative analysis of RhGln-positive punctate structures also confirmed that Cx43 HC-mediated transfer of glutamine from astrocytes to synapses occurs under basal conditions and is significantly enhanced by synaptic stimulation (Fig. 5a-c). Remarkably, restoring *in vivo* Cx43 expression selectively in hippocampal astrocytes from -/- mice using recombinant adeno-associated viruses (rAAV) (Fig. 6a-c) fully rescued activity-dependent RhGln transfer (-/- Cx43 rescue), while the GFP control virus (-/- GFP Ct) had no effect (Fig. 6d-g), further supporting a direct functional role of Cx43 in astroglial glutamine transfer.

**Fig. 5.**
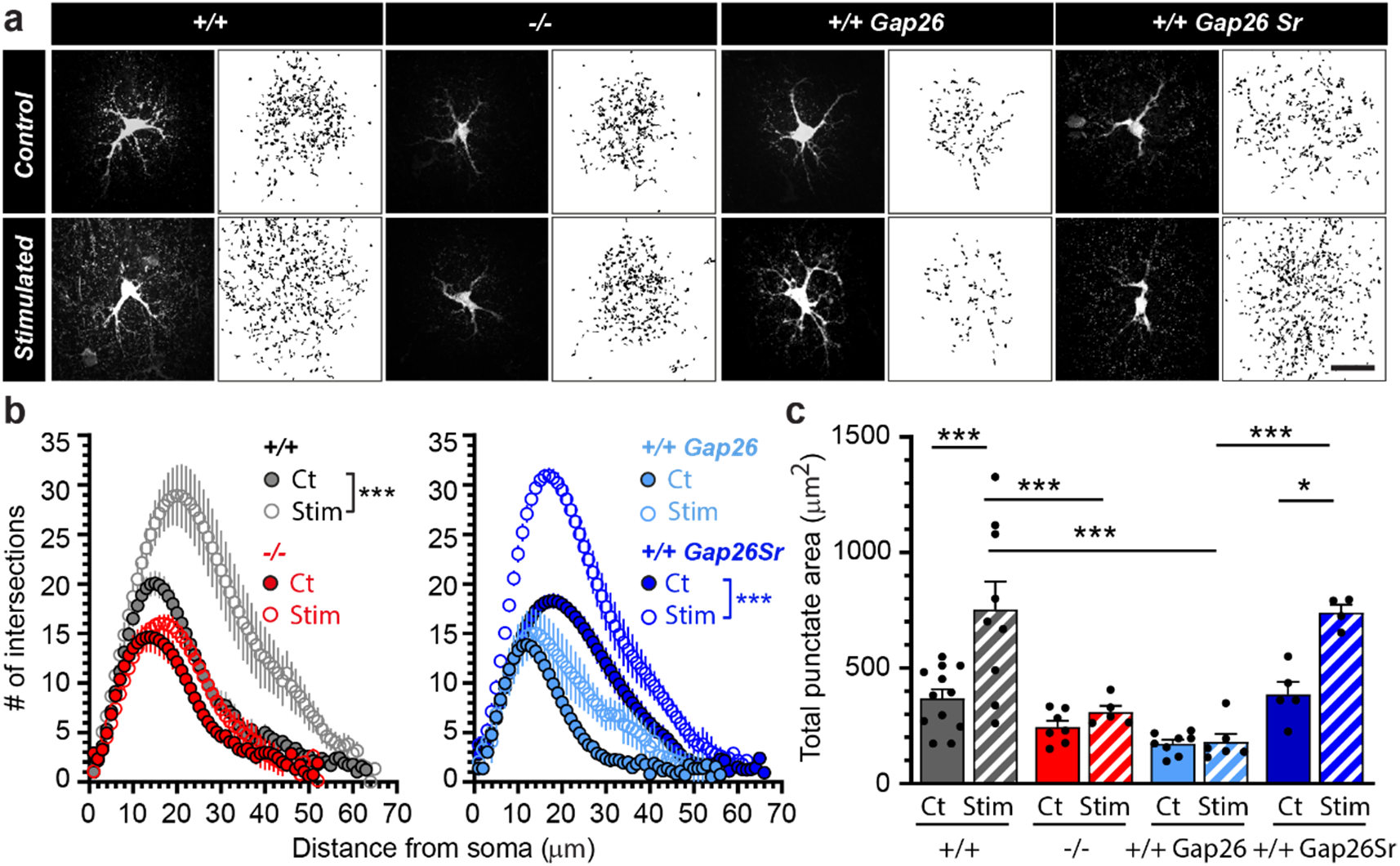
Cx43 hemichannels mediate activity-dependent transfer of glutamine from astrocytes to synapses. **a**, Representative confocal (dark background) and thresholded binary (white background) images of hippocampal CA1 astrocytes dialyzed with RhGln (0.8mM, 20min) via the patch pipette under control or stimulated (10 Hz, 30s every 3min for 20min) conditions in acute slices obtained from: wild type mice (+/+), glial conditional Cx43 knockout mice (-/-) or wild type mice exposed to Gap26 (+/+ Gap26) or Gap26 scramble (+/+ Gap26Sr) peptides. The binary images were quantified by Sholl analysis **(b)** and total punctate area **(c)**. In +/+ mice, repetitive synaptic stimulation strongly increased the punctate RhGln-labeling compared to control as shown by both Sholl analysis (+/+: Ct, n=12; Stim, n=9, *p*<0.0001, Two-way ANOVA in **b**) and total punctate area (*p*<0.0001 between Ct and Stim in +/+, One-way ANOVA with Bonferroni’s post hoc test in **c**). This was abolished in -/- mice (-/-: Ct, n=7; Stim, n=5, *p*=0.1097 in **b**; *p*>0.999 between Ct and Stim in -/-, and *p*=0.0002 between Stim of +/+ and -/- in **c**) or in the presence of Gap26 (+/+ Gap26: Ct, n=8; Stim, n=5, *p*=0.3044 in **b;** *p*>0.999 between Ct and Stim in +/+ Gap26, and *p*<0.0001 between Stim of +/+ and +/+ Gap26 in **c**), but unchanged in the presence of the scramble Gap26 peptide (+/+ Gap26Sr: Ct, n=5; Stim, n=4, *p*<0.0001 in **b**; *p*=0.0143 between Ct and Stim with +/+ Gap26Sr, and *p*<0.0001 between Stim of +/+ Gap26 and +/+ Gap26Sr in **c**); Two-way ANOVA was used in **b** and One-way ANOVA with Bonferroni’s post hoc test in **c** Scale bar: **a,** 20μm. Asterisks indicate statistical significance (***: *p*<0.001; *: *p*<0.05).

**Fig. 6.**
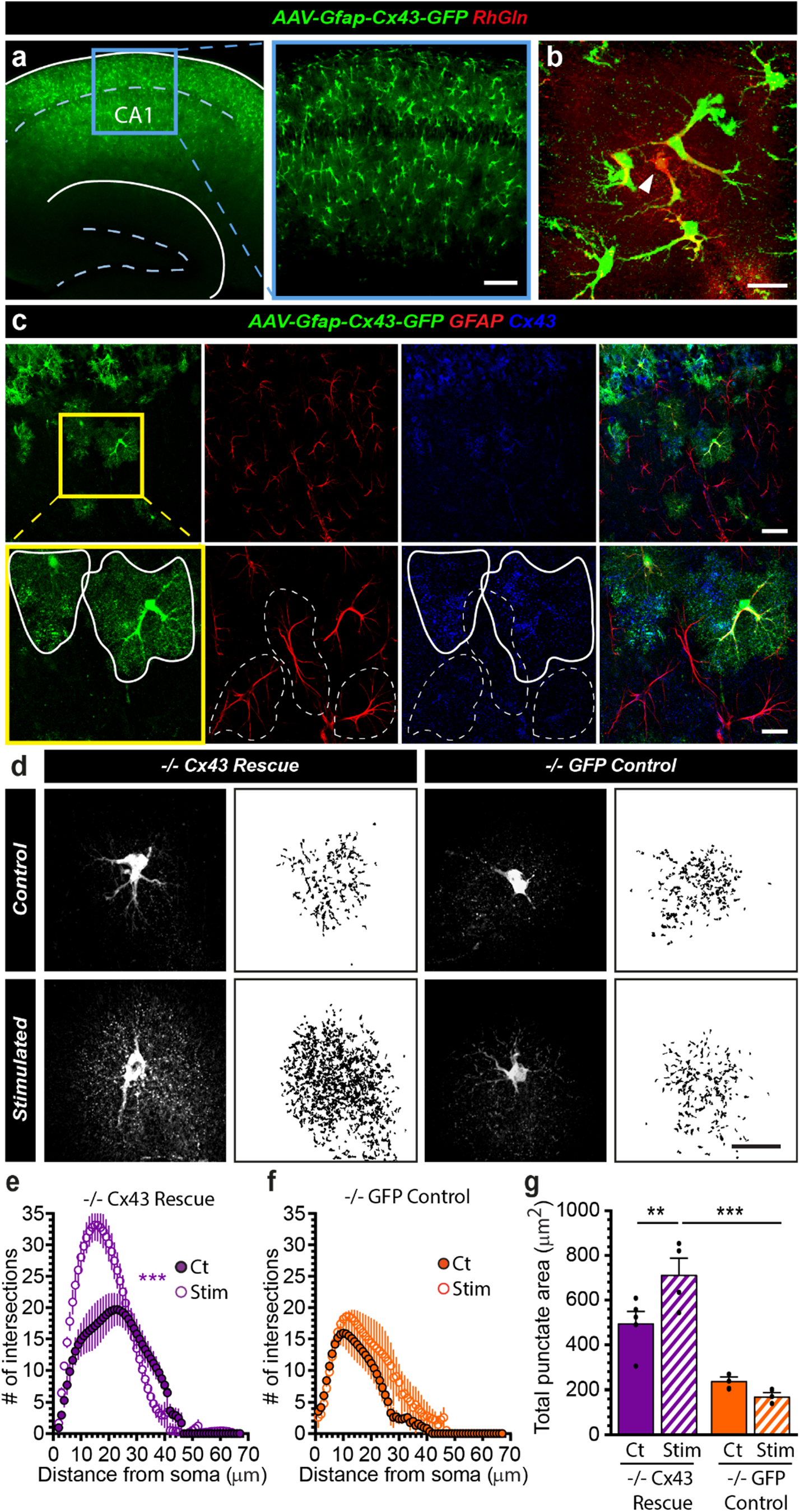
Restoring *in vivo* Cx43 expression in hippocampal astrocytes from Cx43-/- mice rescues activity-dependent transfer of glutamine. **a**, Sample image of the hippocampus of Cx43-/- mice injected intra-hippocampally with rAAV2/9-GFAP-Cx43-GFP virus, showing numerous cells expressing Cx43-GFP in the CA1 area. The blue box is magnified on the right. **b,** RhGln (red) was loaded into a Cx43-GFP expressing astrocyte (green) via a patch pipette as shown. Arrow head indicates the patched cell. **c,** Sample images after immunostaining showing specific expression of Cx43 (blue) in Cx43-GFP-positive (green) astrocytes (GFAP, red). The yellow box is magnified in the bottom row. Solid and dotted white lines outline GFP-positive and -negative astrocytes, respectively. **d-g,** Cx43-/- mice first received either rAAV2/9-GFAP-Cx43-GFP (-/- Cx43 Rescue, **d left, e and g**) or rAAV2/9-GFAP-GFP (-/- GFP Control, **d right, f and g**) virus. Hippocampal astrocytes were then dialyzed with RhGln under either control or synaptic stimulation (10Hz, 30s) conditions for 20min. Both representative confocal (dark background) and thresholded binary (white background) images are shown in **d** for each condition. The binary images were quantified by Sholl analysis **(e and f)** and total punctate area **(g)**. The stimulation-induced transfer of RhGln was rescued in Cx43-/- mice (n=5) by restoring Cx43 expression selectively in astrocytes via viral infection shown by both Sholl analysis (-/- Cx43 rescue, n=4, *p*<0.0001, Two-way ANOVA for **e**) and total punctate area (*p*=0.0042 between Ct and Stim with -/- Cx43 Rescue, One-way ANOVA with Bonferroni’s post hoc test for **g**) as compared to the GFP control infection (-/- GFP Ct, n=3; Stim, n=3, *p*=0.9856, Two-way ANOVA for **f**, *p*>0.999 between Ct and Stim with -/-GFP Control and *p*<0.0001 between Stim of -/- Cx43 Rescue and -/- GFP Control, One-way ANOVA with Bonferroni’s post hoc test for **g**). Scale bars: **a,** 200μm (left) and 50μm (right); **b,** 50μm (top) and 20μm (bottom); **c-d,** 20μm. Asterisks indicate statistical significance (***: *p*<0.001; **: *p*<0.01).

### Astroglial glutamine supply via Cx43 hemichannels is required for synaptic transmission and recognition memory

With direct visual evidence of an activity- and Cx43-dependent mobilization of astroglial glutamine into synaptic structures, we next investigated the functional relevance of this process. To determine its contribution to the replenishment of the presynaptic pool of glutamate, we examined fEPSPs at hippocampal CA3 to CA1 synapses in response to 10Hz repetitive stimulation of Schaffer collaterals in acute slices (Fig. 7a-b). Upon stimulation at 10Hz, a fast synaptic facilitation followed by a depression, resulting from presynaptic glutamate depletion, was observed (Fig. 7b-d). We found that, compared to control +/+ slices, excitatory synaptic transmission depressed faster in -/- slices as well as in +/+ slices treated with Gap26, but not with Gap26Sr (Fig. 7c-e). This indicates that Cx43 HCs are crucial for sustaining repetitive excitatory synaptic activity. We then tested whether the impairment of excitatory synaptic transmission upon loss of Cx43 resulted from lack of an astroglial glutamine supply. To do so, we treated acute hippocampal slices with exogenous glutamine (4mM for 1-4hr), and found in these slices that Cx43 genetic disruption in astrocytes (-/-) or acute pharmacological Gap26 treatment no longer impaired excitatory transmission upon repetitive stimulation (Fig. 7f-g). Thus, exogenous glutamine fully rescued synaptic activity to wild type levels. Importantly, the same exogenous glutamine treatment in +/+ slices had no effect on excitatory transmission (Fig. 7h), indicating that endogenous glutamine level in +/+ slices is sufficient to sustain synaptic activity under physiological conditions. Altogether, these findings argue that Cx43 HCs contribute to physiological excitatory synaptic transmission by fueling synapses with glutamine. The hippocampus is well-known for its functions in learning and memory, including recognition memory^26^. Because the presynaptic pool of glutamate contributes to recognition memory^27^, we investigated *in vivo* the functional significance of Cx43 HC-mediated release of glutamine in such form of hippocampal memory. To do so, we administered intra-hippocampal injections of Gap26 or control Gap26Sr to adult wild type mice, then subjected them to a novel object recognition test (Fig. 7i). As mice are naturally attracted towards novelty, recognition memory was assessed by the relative time spent exploring a novel versus familiar object. While control Gap26Sr-injected mice spent significantly more time exploring the novel object, this preference was not present in Gap26-treated mice, indicating impaired recognition memory (Fig. 7j). Remarkably, we found that mice co-injected with Gap26 and glutamine no longer exhibited memory impairment, implying that exogenous glutamine fully rescued object recognition (Fig. 7j), as it did for synaptic transmission (Fig. 7f-g). Our data thus show that the Cx43HC-dependent regulation of hippocampal excitatory synaptic transmission and recognition memory are both mediated by glutamine.

**Fig. 7.**
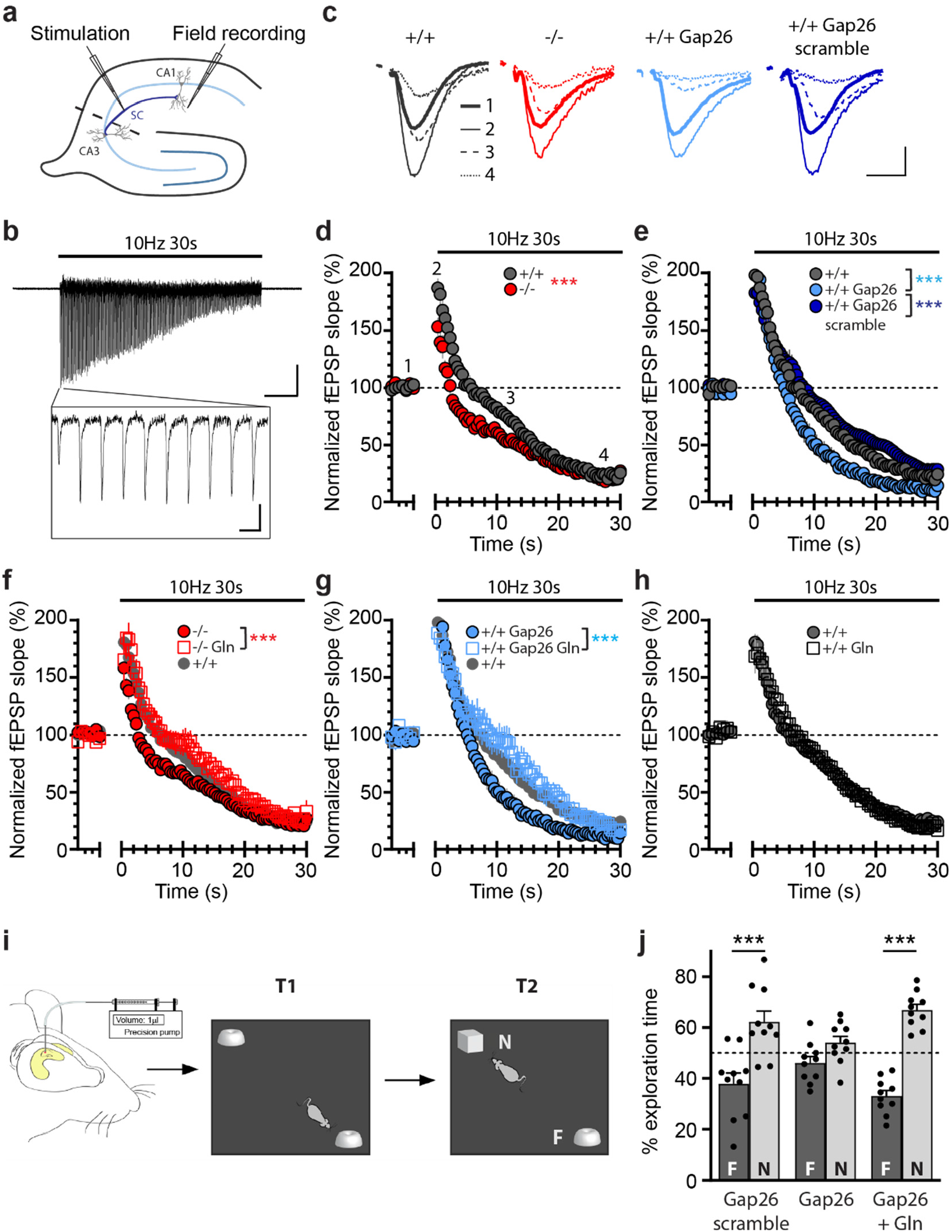
Astroglial glutamine supply via Cx43 hemichannels sustains glutamatergic synaptic transmission and is required for novel object recognition memory. **a,** Schematic drawing depicting recording of field excitatory postsynaptic potentials (fEPSP, field recording) evoked by Schaffer collaterals (SC) stimulation in the CA1 region of hippocampal slices. **b,** Representative trace for fEPSP recorded during stimulation (10Hz, 30s) is presented with the first 10 responses magnified in inset. **c,** Sample traces are shown for fEPSPs recorded before (1, thick solid line), at the start of (2, thin solid line), during (3, thin dashed line), and at the end of (4, thin dotted line) the 10Hz stimulation. **d-e,** Plots of fEPSP slope normalized to baseline upon 10Hz stimulation (horizontal line above) show a decrease in synaptic transmission in slices from -/- (**d**, n=11, *p*<0.0001) or with Gap26 treatment (**e**, +/+ Gap26, n=9, *p*<0.0001) as compared to +/+ (n=11 for **d**, n=14 for **e**), whereas there was no effect of the scramble Gap26 treatment (**e**, +/+ Gap26 scramble, n=6, *p*=0.9994 with +/+, *p*<0.0001 with +/+ Gap26) (Two-way ANOVA). Numbers 1–4 in **d** correspond to regions referred to in **c**. **f-g,** Glutamine pre-treatment (4mM for 4-6h) in -/-slices (**f**, n=16 for -/-, n=8 for -/- Gln, *p*<0.0001) and Gap26 treated slices (**g**, n= 9 for +/+ Gap26, n= 7 for +/+ Gap26 Gln, *p*<0.0001) rescued transmission to +/+ levels (Two-way ANOVA). +/+ plots are also shown for comparison. **h,** Glutamine pre-treatment alone in +/+ mice had no effect (n=16 for +/+, n=8 for +/+ Gln, *p*>0.9999, Two-way ANOVA). **i**, Mice underwent intra-hippocampal injections of either Gap26 scramble (1mM), Gap26 (1mM), or Gap26 (1mM) + Glutamine (200mM) 30min before being submitted to the novel object recognition task which consisted of first an acquisition trial (T1, exploration of 2 similar objects) and then 24h later to a restitution trial (T2, exploration of a familiar object “F” from T1 and a novel object “N”). **j**, Percent exploration time revealed a loss of preference for the novel object in mice treated with Gap26, but was rescued by co-injection of glutamine (Gap26 + Gln). n=10 in each condition (*p*<0.0001 between F and N with Gap26 scramble, *p*=0.2371 between F and N with Gap26, *p*<0.0001 between F and N with Gap26 + Gln, Two-way ANOVA with Bonferroni’s post hoc test, Mean ± SEM). Scale bars: **b**, 0.2mV, 5s (full trace), 100ms (inset); **c,** 0.2mV, 10ms. Asterisks indicate statistical significance (***: *p*<0.001; **: *p*<0.01; *: *p*<0.05).

## Discussion

In this study, we developed a novel fluorescent glutamine molecule to directly track glutamine in live cells. Using this probe, we provide direct evidence for activity-dependent redistribution and supply of glutamine from perisynaptic astrocytes to synaptic structures under physiological conditions. Furthermore, we identified astroglial Cx43 HCs as the key mediators of this process, essential for sustaining physiological excitatory synaptic transmission as well as recognition memory. Our findings thus address a long-standing controversy and directly demonstrate a requirement for astroglial glutamine in supporting both physiological synaptic transmission and cognition via an entirely novel mechanism mediated by Cx43 HCs.

### A novel glutamine fluorescent probe to assess astroglial glutamine supply in situ in live cells

Up to now, the occurrence and role of astroglial glutamine supply during physiological processes has lacked support from tools that can directly evaluate glutamine cellular transfer. Initially, radiolabeled glutamine and electron microscopy of glutamine immunogold labeling enabled quantification of brain glutamine levels^16,17^ and glutamine subcellular localization^28^, respectively. However these approaches cannot provide dynamic information about changes in glutamine content. More recently, FRET-based biosensors have been reported to dynamically monitor intracellular levels glutamine^18,19^. However, these genetically-introduced sensors lack the ability to provide spatially-resolved directionality of glutamine flow between different cell types. Similar to the use of a fluorescently tagged glucose^29^ designed to probe local and fast brain metabolism, here we have taken a direct approach to track cellular glutamine transfer in live cells *in situ* using fluorescently tagged glutamine molecules.

Our probe, RhGln, exhibits several key features required for live detection of glutamine in cellular compartments. Its water solubility permits use in physiological extracellular and intracellular milieu. Its bright fluorescence allows clear detection in small processes using super resolution imaging. Its small size of 0.64 kDa enables intracellular and extracellular diffusion, including through small transmembrane channels such as Cx HCs. Finally, its stability as a non-hydrolysable glutamine analog enables detection of glutamine rather than its metabolites. We have indeed engineered a fluorescent tag on the amide side chain, preventing conversion of glutamine to glutamate. As glutamine is readily metabolized intracellularly, this aspect of our probe is essential to ensuring unambiguous visualization of glutamine mobilization. It also allows us to observe the accumulation of glutamine at its site of storage or conversion.

Using this probe, we show that upon physiological synaptic activity, RhGln that has been selectively delivered to single astrocytes via patch clamp redistributes from the astroglial network into neighboring presynaptic structures. We tracked the probe at a subcellular level, revealing activity-dependent punctate labeling surrounding the RhGln-injected astrocyte, that was not associated with changes in astroglial morphology, but was localized to presynaptic structures, as revealed by super resolution STED microscopy. This finding demonstrates the occurrence of a directional glutamine cellular transfer from astrocytes to presynaptic elements under physiological conditions. Of note, the specific RhGln spatial pattern displayed upon activity, consisting of punctate labeling of presynaptic compartments, was not observed with unconjugated Rh dye alone, and was inhibited upon blockade of neuronal glutamine transport. This indicates that upon activity, RhGln enters synaptic compartments via neuronal glutamine transport, in contrast to the inert control dye, and thus retains the specificity of glutamine cellular trafficking. Our probe therefore offers an advantage over existing methods and is uniquely suitable for investigating directional transfer of glutamine in live cells. Its use in hippocampal tissues here provides the first and direct visual evidence of an activity-dependent mobilization of astroglial glutamine into presynaptic terminals in physiological conditions *in situ*.

### Astroglial glutamine supply sustains physiological activity

The pool of presynaptic glutamate is limited. According to estimates, this pool would be exhausted within minutes of physiological synaptic activity without replenishment. This estimate held true at both small presynaptic terminals from hippocampal CA3 to CA1 synapse and larger synapses at the calyx of Held synapse^6^. The mechanism by which astrocytes could locally replenish the presynaptic pool of glutamate has thus been seen as a suitable pathway to sustain brain activity in the absence of glutamine synthesis at excitatory synapses^4,5^. Astrocytes are indeed a well-established source of recycled glutamate. Their numerous perisynaptic processes, strongly enriched in glutamate transporters^2,30^, provide efficient glutamate clearance upon synaptic activity, while their specific expression of glutamine synthetase enables the conversion of sequestered glutamate into glutamine^3^. Surprisingly, despite the need of rapid glutamate recycling for normal neurotransmission, until now the significance of the glutamate-glutamine cycle has only been clearly evident during high glutamate turnover, for example epileptiform activity^7–9^. The occurrence and relevance of this cycle under physiological conditions remains controversial, not only due to lack of tools to directly track glutamine transfer, but also due to conflicting findings on neuronal activity. Previous work in cortical and hippocampal neurons *in vitro*^11,31^, hippocampal slices^10^, retina^12^, and *in vivo*^13^ suggested that the glutamate-glutamine cycle is not required for physiological synaptic transmission, network activity or memory. In contrast, other studies performed in brainstem slices^14^, isolated excitatory terminals from hippocampal slices or active neocortical slices^15^ reported a role for presynaptic glutamine transport and glutamine synthesis, respectively, in supporting glutamatergic synaptic transmission. These contrasting findings likely stem from different experimental paradigms, including the use of distinct tools to interfere with the cycle, variations in preparations, brain regions and types of synapses, which are not directly comparable due to disparate presynaptic pool size, release probability, and thus variation in glutamine requirement. Furthermore, these studies mostly relied on pharmacological blockers to perturb different components of the cycle, and may thus suffer from incomplete inhibition as well as significant off-target effects.

Here, using a novel multidisciplinary approach combining chemical biology to synthetize a fluorescent glutamine probe, super resolution imaging, electrophysiology and behavioral testing, we demonstrate a requirement for astroglial glutamine in sustaining physiological synaptic and cognitive demands. We uncovered a significant glutamine-dependent component of excitatory transmission evoked by synaptic stimulation at 10Hz, a frequency within the physiological range for hippocampal CA3 to CA1 synapses^32^, known to initially facilitate glutamate release and then to deplete presynaptic pools^33^. This glutamine component was essential not only for the initial facilitation of excitatory synaptic transmission, but also for its maintenance, indicating its crucial role in the replenishment of the presynaptic pool of glutamate. We showed that astrocytes are the source of glutamine, and supply is dependent on Cx43, a gap junction channel subunit abundantly expressed in astrocytes. Using imaging of the RhGln probe, we indeed found that astrocytes, via Cx43, mediate an activity-dependent glutamine supply from astrocytes to synapses. Furthermore, our electrophysiological recordings reveal that excitatory transmission evoked synaptically by the same regime of activity also relies on astrocytes, via Cx43, and glutamine supply. We indeed found that exogenous glutamine fully rescued the impaired excitatory transmission evoked synaptically in mice lacking Cx43 expression conditionally in astrocytes. Importantly, we also found that endogenous levels of glutamine are not only required, but sufficient to sustain evoked excitatory synaptic transmission since exogenously supplying glutamine in wild type mice had no influence on glutamatergic synaptic transmission. Thus, we provide direct quantitative evidence that excitatory synaptic transmission at hippocampal CA3 to CA1 synapses relies on astroglial glutamine.

### Cx43 hemichannels as a novel molecular pathway for astroglial glutamine supply to synapses

Cxs in astrocytes form both GJs and HCs, and underlie the transfer of small molecules within and out of the astroglial network, respectively^34^. GJs, by mediating the extensive network communication of astrocytes, contribute to several key physiological functions, including metabolic support and extracellular homeostasis^34^. In contrast, Cx43 HCs are thought to be active mostly in pathological conditions, where they contribute to pathophysiological processes via release of molecules such as ATP or prostaglandins^35^. Here we reveal that Cx43 HCs are not only active under physiological conditions, but also sustain synaptic transmission and cognition through glutamine release. Although Cx43 GJs have been shown to be permeable to several metabolites such as glucose, lactate or glutamine^36,37^, until now this has not been the case for Cx43 HCs^38^. We thus identify astroglial Cx43 HCs as a novel release pathway for glutamine controlling physiological neuronal activity. Using immunohistochemistry, biochemistry and electron microscopy, we first show that Cx43 is present near hippocampal synapses. We also report functionally that Cx43 HCs open under physiological conditions in an activity-dependent manner. Next, using super resolution imaging and electrophysiology, we found that Cx43 HC activity is essential for the activity-dependent transfer of glutamine from astrocytes to synapses, and that this HC-mediated glutamine supply is crucial for sustaining physiological excitatory synaptic activity. Rescuing Cx43 expression selectively in astrocytes from Cx43 conditional knockout mice indeed restored astroglial glutamine transfer. In addition, blocking Cx43 HC activity, either chronically and non-selectively using astroglial conditional knockout mice, or acutely using specific blocker peptides, impaired astroglial glutamine transfer and excitatory synaptic transmission. We demonstrated that glutamine is required for sustaining synaptic transmission by restoring normal activity with exogenous glutamine. It is noteworthy that a few studies recently reported that Cx43 HCs can regulate neuronal activity via different mechanisms, including synaptic activity and plasticity via release of ATP^23^ or D-serine^39^, or network activity, i.e. olfactory bulb slow oscillations^40^ and hippocampal surround inhibition within synaptically active regions^41^, both via ATP release.

We also report that astroglial glutamine supply via Cx43 HCs is relevant for cognition. We found that glutamine supply associated with Cx43 HC activity is essential for cognitive performance requiring hippocampal memory, as mice with acutely blocked Cx43 HCs, were no longer able to recognize a novel object in the presence of Gap26. This defect was rescued by concomitant *in vivo* delivery of exogenous glutamine. These data are consistent with a recent study indicating a role for Cx43 HCs in spatial short-term memory^42^, although the underlying mechanism was not identified in this work.

It is noteworthy that almost two decades ago it was suggested that multiple pathways are responsible for astroglial glutamine efflux *in vitro*, and that astroglial glutamine transporters only account for ~50% of all glutamine release from astrocytes^43^. The authors attributed the remaining ~50% of basal glutamine efflux to yet unidentified mechanisms, however pharmacologically characterized it as Na^+^-independent. Incidentally, transmembrane channels such as Cx43 GJs can transport metabolites such as glucose, lactate or glutamine^36,37^ in a Na^+^-independent manner. Therefore, our data identifying Cx43 HCs as an entirely novel pathway underlying astroglial glutamine efflux likely accounts for the unknown mechanisms observed in the *in vitro* study. We indeed found that the specific inhibition of Cx43 HC activity also significantly decreased astroglial glutamine transfer by ~50%. Furthermore, activity-dependent increase in glutamine transfer as well as cognitive performance *in vivo*, were nearly completely abolished by knocking out Cx43 or by acute inhibition of Cx43 HC activity. Together with our dye uptake experiments showing that Cx43 HC activity is enhanced by activity, it is likely that Cx43 HC-mediated glutamine release may even account for more than 50% of glutamine efflux upon synaptic activity. An obvious advantage of using Cx43 HCs over specialized transporters to mediate glutamine release from astrocytes is reduced energy cost for the cell. The diffusion of molecules through a large pore along a concentration gradient does not consume ATP, and is thus more energetically favorable than release via transporters.

In summary, we generated and utilized a novel fluorescent probe RhGln to directly observe an activity-dependent transfer of glutamine from astrocyte to hippocampal excitatory synapses and demonstrated a fundamental requirement of the astroglial glutamine supply during physiology. We also showed that this process involves a novel release pathway mediated by Cx43 HCs and confers physiological functional relevance for excitatory synaptic transmission and recognition memory. Our findings thus reveal an essential function for perisynaptic astrocytes in supplying glutamine as a fuel to synapses and has important implications for future studies on astroglial contributions to sustaining physiological brain functions. Because glutamine supply and Cx43 are altered in several neurological diseases, these data also provide an astrocyte basis for cognitive disorders.

## Methods

### Animals

All procedures on animals were performed according to the guidelines of European Community Council Directives of 01/01/2013 (2010/63/EU) and our local animal care committee (Center for Interdisciplinary Research in Biology in College de France). Experiments were carried out using mice of wild type C57BL/6j background, mice expressing enhanced green fluorescent protein under the astrocytic promoters glial fibrillary acidic protein (GFAP-eGFP) or aldehyde dehydrogenase 1 family member L1 (Aldh1l1-eGFP), as well as mice with glial conditional deletion of connexin 43 (Cx43) in astrocytes Cx43^fl/fl^:hGFAP-Cre (Cx43-/-). GFAP-eGFP and Cx43-/- mice were provided by F. Kirchhoff (University of Saarland, Germany) and K. Willeke (University of Bonn, Germany), respectively, and were previously characterized^44,45^. Bac Aldh1l1-eGFP mice were obtained from The Jackson Laboratory (GENSAT project; Stock No: 030247) and were characterized^46^. Mice of both genders and littermates were used at postnatal days (P) 17-30 or 3 months of age. All efforts were made to minimize the number of animals used and their suffering.

### Recombinant adeno-associated virus (rAAV) generation and stereotaxic surgery

For rAAV *in vivo* gene transfer, a transgene composed of GFP and Cx43 cDNA separated by a P2A sequence in a single open reading frame was placed under the control of a GFAP-specific promoter in a rAAV shuttle plasmid containing the inverted terminal repeats (ITR) of AAV2 (AAV-GFAP-Cx43-GFP). Pseudotyped serotype 9 rAAV particles were produced by transient co-transfection of HEK-293T cells, as previously described^47^. Viral titers were determined by quantitative PCR amplification of the ITR on DNase-resistant particles and expressed as vector genome per ml (vg/ml). Cx43-/- mice (P15-17) were deeply anesthetized using mixture of ketamine (95mg/kg; Merial) and xylazine (10mg/kg; Bayer) in 0.9% NaCl and fitted into a stereotaxic frame (David Kopf Instruments). 0.5μl of either rAAV2/9-GFAP-Cx43-GFP or rAAV2/9-GFAP-GFP (1.5 x 10^12^vg/ml) was unilaterally injected into the hippocampus at the rate of 0.1μl/min using the following coordinates from Bregma: antero-posterior −1.95mm; medio-lateral −1.5mm; dorso-ventral −1.38mm. The injection was performed using 29-gauge blunt-tip needle connected to 2μl Hamilton syringe and the injection rate was controlled by syringe pump (KD Scientific). After the injection, the needle was left in place for 5 min and then slowly withdrawn. Following surgery, mice were allowed to recover from anesthesia on a heating pad and monitored for the next 24h. Mice were sacrificed 13-15 days post-surgery for electrophysiology or immunohistochemistry experiments.

### Antibodies, immunohistochemistry and immunoblotting

All antibodies were commercially available and validated by manufacturers. The following primary antibodies were used: polyclonal chicken anti-GFP (1:500, AB13970, Abcam), monoclonal mouse anti-VGlut1 (1:200, 135511, Synaptic Systems), monoclonal mouse anti-Cx43 (1:500, 610-062, BD Biosciences), polyclonal rabbit anti-Cx43 (1:500, 71-2200, Zymed Laboratories), monoclonal mouse anti-GFAP (1:500, G3893, Sigma), and monoclonal mouse anti-GAPDH-peroxidase (1:1000, G9295, Sigma). The following secondary antibodies were used for immunohistochemistry: goat anti-mouse IgG conjugated to Alexa 488, 555 or 647 (1:2000; A11029, A21424 or A21235, Life Technologies), goat anti-rabbit IgG conjugated to Alexa 488 or 555 (1:2000, A11034 or A21429, Life Technologies), and goat anti-chicken IgG conjugated to Alexa 488 (1:2000, A11039, Life Technologies). The following secondary antibodies were used for immunoblotting: goat anti-mouse IgG-HRP (1:2500, sc-2005, Santa-Cruz) and goat anti-rabbit IgG-HRP (1:2500, sc-2004, Santa-Cruz). Immunohistochemistry and immunoblotting were performed as previously described^48,49^. Briefly, acute hippocampal slices (300-400μm) were fixed in 4% paraformaldehyde (PFA) at room temperature for 1h then blocked for 1h in phosphate buffered saline (PBS) containing 1% Triton-X100 and 1% gelatin. Slices were then incubated in appropriate primary antibodies at 4°C overnight followed by secondary antibodies at room temperature for 2h the next day. Sections were mounted in Fluoromount-G™ (Thermo Fisher, USA) for image acquisition. Western immunoblotting was carried out by separating equal amounts of protein by electrophoresis in a 10% polyacrylamide gel. This was followed by transfer of proteins onto nitrocellulose membranes which were then saturated with 5% fat-free dried milk in triphosphate buffer solution and incubated overnight at 4°C with appropriate primary antibodies. On the next day, membranes were treated with peroxidase-conjugated secondary antibodies for 2h at room temperature and revealed using a chemiluminescence detection kit (ECL, GE Healthcare).

### Synthesis, structural confirmation and characterization of rhodamine-tagged glutamine

The red fluorescent molecule rhodamine provides high molar absorptivity, fluorescent quantum yield and photostability and was therefore selected as the fluorescent tag of choice for glutamine. In addition, having a probe in the red spectrum allows it to be used in transgenic mice expressing green fluorescent protein under specific promotors. For this study, rhodamine tagged-glutamine (RhGln) molecule was synthesized in a 5-step chemical reaction detailed in Supplementary Fig. 1. For structural confirmation of the intermediates compounds 3 and 4 and final product RhGln, proton and carbon-13 nuclear magnetic resonance (1H and 13C NMR) spectrometry as well as mass spectrometry were used (Supplementary Fig. 2). The 1H and 13C spectra were collected on a JEOL-FT NMR 400 MHz (100 MHz for 13C NMR) spectrophotometer using CDCl3 or CD3OD as a solvent and tetramethylsilane as an internal standard. Mass spectrometry analyses were performed by Small Molecule Mass Spectrometry platform of ICSN (Centre de Recherche de Gif-sur-Yvette, Universite Paris-Saclay, France). All chemicals and reagents were purchased from Sigma-Aldrich and used without any further purification. **4-(2-phthalimidoethoxy) rhodamine (compound 3):** A solution of 1 (0.31g, 1.06mmol) in propionic acid (5mL) was stirred at room temperature followed by the addition of 8-hydroxyjulolidine (0.40g; 2.13mmol) and PTSA (*p*-toluenesulfonic acid; catalytic amount). The resulting reaction mixture was protected from light and stirred at room temperature for 12h. Saturated sodium acetate solution was added to the reaction mixture until pH reached 7-8, resulting in the precipitation of intermediate compound 2. The precipitate was filtered, washed with water, dried and used immediately for the next step. The crude product 2 (0.60g; 0.91mmol) was dissolved in dry CH_2_Cl_2_ (20 mL) followed by the addition of chloranil (0.33g; 1.37mmol) and allowed to stir for 12h at room temperature. The solvent was removed under reduced pressure and the residue was subjected to column chromatography (CH_2_Cl_2_/MeOH; 9.5/0.5). Compound 3 was obtained as a purple solid (79% over 2 steps). ^1^H NMR (CDCl_3_, 400 MHz): *δ* = 7.87-7.89 (m, 2 H, Ar-H), 7.74-7.76 (m, 2 H, Ar-H), 7.21 (d, *J* = 8 Hz, 2 H, Ar-H), 7.08 (d, *J* = 8 Hz, 2 H, Ar-H), 6.82 (s, 2 H, Ar-H), 4.35 (t, *J* = 8 Hz, 2 H, CH_2_), 4.18 (t, *J* = 8 Hz, 2 H, CH_2_), 3.51-3.58 (m, 8 H, CH_2_), 3.02 (t, *J* = 8 Hz, 4 H, CH_2_), 2.69 (t, *J* = 8 Hz, 4 H, CH_2_), 2.10 (t, *J* = 4 Hz, 4 H, CH_2_) and 1.97 (t, *J* = 4 Hz, 4 H, CH_2_) ppm. ^13^C NMR (CDCl_3_, 100 MHz): *δ* = 168.31, 159.55, 154.72, 154.66, 152.31, 151.11, 134.35, 132.04, 131.09, 126.81, 125.17, 123.57, 114.96, 113.00, 105.50, 65.01, 51.04, 50.59, 37.28, 27.78, 20.77, 20.02 and 19.84 ppm. HRMS (TOF MS ES+): found: [M]^+^ = 636.2865; Calc. for C_41_H_38_N_3_O_4_^+^: [M]^+^ = 636.2857. **4-(2-aminoethoxy) rhodamine (compound 4):** To a solution of compound 3 (0.12g, 0.19mmol) in absolute ethanol (5mL) was added hydrazine monohydrate (0.09g, 1.95mmol). The resulting mixture was refluxed for 4h. The solvent was removed under reduced pressure and 20ml of CH_2_Cl_2_ was added to the mixture. The organic layer was washed with 10ml of NaOH solution (10^−4^ mol L^−1^) and was dried with MgSO_4_ to provide 70mg of free amine 4 (82%), which was used without any further purification. ^1^H NMR (CD3OD, 400 MHz): *δ* = 7.33 (d, *J* = 8 Hz, 2 H, Ar-H), 7.21 (d, *J* = 8 Hz, 2 H, Ar-H), 6.94 (s, 2 H, Ar-H), 4.21 (t, *J* = 4 Hz, 2 H, CH_2_), 3.70 (t, *J* = 4 Hz, 2 H, CH_2_), 3.50-3.56 (m, 8 H, CH_2_), 3.06 (t, *J* = 8 Hz, 4 H, CH_2_), 2.72 (t, *J* = 8 Hz, 4 H, CH_2_), 2.08-2.11 (m, 4 H, CH_2_) and 1.95-1.96 (m, 4 H, CH_2_) ppm. ^13^C NMR (CD3OD, 100 MHz): *δ* = 160.17, 154.90, 152.31, 151.02, 131.08, 126.46, 125.02, 123.79, 114.53, 112.66, 105.27, 50.48, 49.98, 40.16, 29.42, 27.29, 20.48, 19.63 and 19.53 ppm. HRMS (TOF MS ES+): found: [M]^+^ = 506.2811; Calc. for C_33_H_36_NSO_2_^+^: [M]^+^ = 506.2802. **4-(2-(N-glutamino)ethoxy) rhodamine (RhGln):** Compound 4 (0.10g, 0.21mmol), Boc-Glu-OtBu (0.06g, 0.21mmol), EDCI.HCl (0.04g, 0.23mmol) and DMAP (0.002g, 0.02mmol) were dissolved in dry DMF (5mL) under argon atmosphere. The mixture was stirred at room temperature for 12h followed by solvent removal under reduced pressure. The residue was dissolved in CH_2_Cl_2_ (50mL) and washed with water (100mL). The organic layer was dried over Na_2_SO_4_ and concentrated under vacuum. The residue was then dissolved in a mixture of CH_2_Cl_2_-TFA (10mL; 8:2 v/v) and the resulting mixture was allowed to stir at room temperature for 1h. The solvent was removed under reduced pressure and the remaining solid was subjected to column chromatography (CH_2_Cl_2_/MeOH; 9/1). RhGln was obtained as a purple solid (60% over 2 steps). ^1^H NMR (CD3OD, 400 MHz): *δ* = 7.34 (d, *J* = 8 Hz, 2 H, Ar-H), 7.21 (d, *J* = 8 Hz, 2 H, Ar-H), 6.94 (s, 2 H, Ar-H), 4.19 (t, *J* = 8 Hz, 2 H, CH_2_), 3.75-3.78 (m, 1 H, CH), 3.67 (t, *J* = 4 Hz, 2 H, CH_2_), 3.51-3.57 (m, 8H, CH_2_), 3.07 (t, *J* = 8 Hz, 4 H, CH_2_), 2.88-2.91 (m, 2 H, CH_2_), 2.73 (t, *J* = 8 Hz, 4 H, CH_2_), 2.52 (t, *J* = 4 Hz, 2 H, CH_2_), 2.09-2.17 (m, 6 H, NH_2_ and CH_2_) and 1.97-1.98 (m, 4 H, CH_2_) ppm. ^13^C NMR (CD3OD, 100 MHz): *δ* = 173.84, 164.99, 160.15, 154.98, 152.35, 151.05, 131.05, 126.47, 123.81, 114.49, 112.68, 105.28, 66.45, 53.79, 50.49, 49.99 42.20, 38.78, 31.47, 27.30, 26.43, 20.49, 19.63 and 19.54 ppm. HRMS (TOF MS ES+): found: [M]^+^ = 635.3220; Calc. for C_38_H_43_N_4_O_5_+ [M]^+^ = 635.3228.

The fluorescence properties of RhGln were characterized using both steady state and time resolved measurements. For steady state measurements, samples were prepared in 1cm^2^ quartz optical cell using the appropriate solvent (i.e. chloroform, ethanol or intracellular solution). UV/Vis absorption spectra were recorded on a Varian Cary 5000 spectrophotometer and corrected emission spectra were collected on a Jobin-Yvon SPEX Fluoromax-4 spectrofluorometer using samples with optical densities below 0.1 at the excitation wavelength, as well as at all emission wavelengths. The RhGln exhibits a strong S_0_ → S_1_ transition in the green-to-red region of the visible spectra with a corresponding molar absorptivity of l.2×10^5^M^−1^.cm^1^ at 580nm. The fluorescence quantum yields were determined using Rhodamine 101 in methanol as standard^50^. For time resolved measurements, fluorescence emission decays were obtained using a time correlated single-photon counting (TCSPC) apparatus. Samples were prepared in 1cm^2^ quartz optical cell using the appropriate solvent (i.e. chloroform, ethanol or intracellular solution). Excitation was achieved using a Spectra-Physics setup composed of a Ti:Sa Tsunami laser (FWHM = 100fs) pumped by a frequency doubled Millennia Nd:YVO4 laser. Light pulses were selected by optoacoustic crystals at a repetition rate of 4MHz and a frequency doubling crystal is used to reach the desired excitation wavelength of 495nm (FWHM = 200fs; 200μW). Emitted photons were detected, through a monochromator centered at 600nm, using a Hamamatsu MCP R3809U photomultiplier connected to a constant-fraction discriminator. The time-to-amplitude converter was purchased from Tennelec. The fluorescence data were analyzed by a nonlinear least-squares method on Globals software developed at the Laboratory of Fluorescence Dynamics at the University of Illinois, Urbana-Champaign. Satisfactory fits of time-resolved fluorescence measurements in intracellular solution were obtained by considering a bi-exponential model, leading to corresponding lifetimes of: τ_1_ = 4.2ns (a_1_ = 97%) and τ_2_ = 0.9ns (a_2_ = 3%). In contrast, single exponential behavior, with nearly identical lifetime (τ = 3.5ns), was observed in non-polar organic solvent such as chloroform. Such effect could derive from the presence of different conformers in intracellular solution, where efficient photo-induced electron transfer could occur between the dye and the glutamine moiety.

### Acute hippocampal slice preparation

Acute transverse hippocampal slices (300-400μm) were prepared as previously described^51^. Briefly, mice were sacrificed by cervical dislocation and decapitation. The hippocampi were immediately isolated and sectioned at 4°C using a vibratome (Leica) in an artificial cerebrospinal fluid (ACSF) containing (in mM): 119 NaCl, 2.5 KCl, 2.5 CaCl_2_, 1.3 MgSO_4_, 1 NaH_2_PO_4_, 26.2 NaHCO_3_ and 11 glucose (pH = 7.4). Slices were maintained at room temperature in a storage chamber containing ACSF saturated with 95% O_2_ and 5% CO_2_ for at least 1h prior to experiments.

### Electrophysiology and dye-filling of astrocytes in acute hippocampal slices

Acute hippocampal slices were transferred to a submerged recording chamber mounted on an Olympus BX51WI microscope and were perfused with ACSF at a rate of 2ml/min at room temperature. Except when indicated, all electrophysiological recordings were performed in the presence of picrotoxin (100μM) and 3-((*R*)-2-carboxypiperazin-4-yl)-propyl-1-phosphonic acid (CPP; 10μM; Tocris Bioscience), and a cut was made between CA3 and CA1 regions to prevent the propagation of epileptiform activity. Neuronal field recordings were performed as previously described^48^. Field excitatory postsynaptic potentials (fEPSPs) were evoked by Schaffer collaterals stimulation (10-20μA, 0.1ms) through an ACSF-filled glass pipette located at a distance of 200μm from the recording area. Basal activity was measured at 0.1Hz whereas enhanced stimulated activity was recorded at 10Hz for 30s once or repeated every 3min for 20 min. For patch-clamp recordings and dye-filling experiments, CA1 astrocytes were visually identified and filled under whole-cell configuration via glass pipettes (5-8MΩ) containing an intracellular solution (in mM): 105 K-gluconate, 30 KCl, 10 HEPES, 10 phosphocreatine, 4 ATP-Mg, 0.3 GTP-Tris, 0.3 EGTA (pH 7.4, 280mOsm/l). Stratum radiatum astrocytes were verified by their low input resistance (~20MΩ), high resting potentials (~-80mV) and linear passive input/output relationship under voltage clamp configuration. 0.8mM of either RhGln or a rhodamine only dye (Rh) was also included for dye-filling experiments and astrocytes were dialyzed for 20min. In some experiments, pharmacological substances were included in the perfusion ACSF before and during the experiments: a Cx43 hemichannel blocker peptide and its scramble version (Gap26/Gap26 scramble; 100μM) and a blocker of neuronal glutamine transporter, α-(Methylamino) isobutyric acid (MeAIB, 20mM) with 5min pre-incubation; a gap-junction blocker carbenoxelone (CBX, 200μM) with 10min pre-incubation; glutamine (4mM) with 1-4h pre-incubation in storage chamber). After dye filling, patch pipettes were carefully removed and slices were fixed for 1h in 4% PFA at room temperature and preserved for immunohistochemistry or directly mounted for image acquisition. Recordings were acquired with Axopatch-1D amplifiers (Molecular Devices, USA), digitized at 10kHz, filtered at 2kHz, stored and analyzed on a computer using pClamp9 and Clampfit10 software (Molecular Devices). fEPSP slopes were normalized to baseline and plotted over 30s during stimulation. Stimulation artifacts were removed in representative traces for illustration.

### Ethidium bromide uptake assay and RhGln bulk loading experiments

Acute hippocampal slices (350μm) were transferred to a submerged recording chamber and incubated in HEPES buffer containing (in mM): 140 NaCl, 5.5 KCl, 1 MgCl_2_, 1.8 CaCl_2_, 10 HEPES, 10 C_6_H_12_O_6_ (pH = 7.4). Ethidium bromide (EtBr; 314 Da, 4μM) or RhGln (0.05mM) was then added extracellularly to the recording chamber. After a 10min pre-incubation, some slices were electrically stimulated (10Hz for 30s every 3min) at Schaffer collaterals for 20min via a glass pipette as described for electrophysiological experiments. To investigate the contribution of Cx43 hemichannels to EtBr uptake, slices were pre-treated 5min before and during EtBr incubation with a Gap26 or Gap26 scramble (100μM). Slices were then rinsed for 15min in fresh HEPES buffer, fixed for 1h in 4% PFA at room temperature and preserved for immunohistochemistry or directly mounted for image acquisition.

### Isolation of synaptosomal fraction

Crude synaptosomal fractions from hippocampus were isolated for immunoblotting. First, freshly dissected whole hippocampi or acute hippocampal slices were mechanically homogenized using a Potter-Elvehjem homogenizer in ice-cold homogenization buffer (0.32M sucrose, 10mM HEPES, 2mM EDTA, protease inhibitors cocktail; 400μl/hippocampus). Homogenate was then centrifuged at 900 x g for 15min at 4°C. The supernatant (S1) was centrifuged at 16,000 x g for 15min at 4°C. The pellet was then washed and resuspended in fresh ice-cold homogenization buffer (300μl/hippocampus) and centrifuged again at 16,000g for 15min at 4°C. The resulting pellet containing synaptosomes (P2) was resuspended in HEPES lysis buffer (50mM HEPES, 2mM EDTA, protease inhibitors cocktail; 150μl/hippocampus). Samples were briefly sonicated to ensure membrane lysis and used for immunoblots. For normalization, exact protein concentration of each sample was determined using Ionic Detergent Compatibility Reagent (22663, Thermofisher, France).

### Immunogold labeling and serial section electron microscopy

Hippocampal tissue preparation for ultrastructural analysis using serial section electron microscopy was previously described ^48^. Briefly, mice were anaesthetized with Narkodorm^®^ (60mg/kg body weight) and then transcardially perfused with physiological saline followed by an ice-cold 0.1M phosphate-buffered (pH 7.4) fixative containing 4% PFA and 0.1% glutaraldehyde (Polyscience Europe GmbH, Eppelheim, Germany) for 15min. To immunolabel Cx43 with gold particles, 100μm vibratome sections containing the hippocampus were made and treated with a monoclonal mouse anti-Cx43 antibody (1:250, Chemicon Europe, Hampshire, UK) overnight at 4°C followed by a 15nm gold-conjugated anti-mouse secondary antibody (1:40, British Biocell, Cardiff, UK) on the next day for 2h. 10–30 serial ultrathin sections (55 ± 5nm) were cut using a Leica Ultracut S ultramicrotome (Leica Microsystems, Vienna, Austria) and collected on Formvar-coated slot copper grids. Prior to electron microscopic examination, sections were stained with 5% aqueous uranyl acetate for 20min and lead citrate for 5min (Reynolds 1963). Ultrathin sections were inspected and photographed with a Zeiss Libra 120 electron microscope (Fa. Zeiss, Oberkochen, Germany) equipped with a Proscan 2K digital camera and the SIS Analysis software (Olympus Soft Imaging System, Münster, Germany). Individual digital images were further process for final illustrations using the Adobe Photoshop™ and Illustrator™ software packages. For the analysis of average distances between gold particles and the edge of the nearest active zone, 75 synaptic complexes from 8 ultrathin sections were used.

### Confocal image acquisition and analysis

Fixed and mounted acute brain slices were examined using a confocal laser-scanning microscope (TCS SP5, Leica). To avoid high surface background, acquisition was always made 10-30μm below slice surface. Fluorescence images were acquired in sequential mode with different excitation lasers (Argon 488nm, DPSS 561nm, and HeNe 633nm). Three different objectives were used: 20x objective (NA = 0.7) was used to capture single-plane image overviews of the entire CA1 region; 40x objective (NA = 1.25) was used to obtain z-stacks of 1μm-intervals to focus on selected areas in this region; 63x objective (NA = 1.4-0.6) at 2.5 x zoom was used to obtain z-stacks of 0.2μm intervals to closely examine morphology and subcellular compartments. Confocal image analysis was performed using ImageJ software (National Institutes of Health, USA). The size of RhGln-labeled network was determined by counting the number of RhGln-labelled cells in individual experiments and averaging across experiments for each condition. To aid visualization for illustration purposes, RhGln signal in a network of cells were subjected to local contrast normalization using an *Integral Image Filters* plugin and thresholded by intensity. In Fig. 2A, sample images are presented as maximum z-projection. To quantify the extent of RhGln-labeling from the soma of the dye-filled astrocyte, high resolution image stacks (obtained using 63x objective) containing single patched astrocytes were analyzed. A mask of each image of the stacks was generated by thresholding first by intensity and then by size (0.5-2μm^2^) using *Subtract background, Threshold*, and *Analyze Particles* plugins. The same thresholding range was used for images from different experimental conditions to allow direct comparison. The maximum projection of the masks for each astrocyte was then analyzed using *Sholl Analysis* plugin. Results were given as the number of intersections found on each concentric circle from the center of the cell soma at 1μm-radius interval. The fitted curve of the number of intersections over distance for each cell was then extracted and averaged. In addition, total punctate area for each cell was also measured from thresholded image masks. For co-localization analysis between RhGln and VGlut1, fluorescence images from each channel were first separately deconvolved using Huygens software (Scientific Volume Imaging B.V., The Netherlands). Image masks based of intensity thresholding were then generated for each channel and merged to find co-localized areas representing presynaptic structures containing RhGln.

Percentage of co-localization was measured as the co-localized areas normalized to the total RhGln-positive area of the area of interest. Analysis of EtBr uptake was performed on stratum radiatum CA1 GFAP-positive astrocytes. Fluorescence intensity was measured as integrated density in arbitrary units and corrected to background fluorescence. For statistical quantifications, 60 CA1 astrocytes were analyzed per acute slice. For the analysis of astrocytic morphology, image stacks containing single astrocytes immunostained for GFAP were processed by *Sholl Analysis*. After intensity thresholding, images were inverted and the number of intersections of astrocytic processes on each concentric circle from the center of the cell soma at 5μm radius interval. In addition, astrocytic soma and domain sizes were determined by delineating the visible cell body area of Aldh1l1-eGFP signal and entire GFAP positive 2D coverage, respectively.

### Super-resolution stimulated emission depletion (STED) imaging in brain slices

STED imaging was performed to acquire high-resolution images of RhGln-labeled astrocytes in acute brain slices at the level of fine astroglial processes and synaptic structures after immunostaining. Our custom-made upright STED microscope (Abberior Instruments) based on a Scientifica microscope body (Slice Scope, Scientifica) was equipped with an Olympus 100X/1.4NA ULSAPO objective lens. It comprises of a scanner design featuring four mirrors (Quad Scanner, Abberior Instruments) with three excitation lasers at 488nm, 561nm, and 640nm (Abberior Instruments, pulsed at 40/80 MHz). Two additional lasers at 595nm (MPB-C, cw), and 775nm (MPB-C, pulsed at 40/80 MHz) were used to generate STED beams. The conventional laser excitation and STED laser beams were superimposed using a beam-splitter (HC BS R785 lambda/10 PV flat, AHF Analysetechnik). Common excitation power with pulsed excitation ranging from 10–20μW with STED power intensities of up to 200mW in the focal plane were used. Super-resolution images were acquired with pixel size of 30nm, 15-25μm dwell time with 3 times averaging.

### Novel object recognition test

Novel object recognition (NOR) was assessed in 3 months old wild type male mice using a NOR apparatus consisting of a dark grey polypropylene box (40 x 30 x 23cm; length, width, height). A glass rectangle (5 x 5 x 5cm) and a ceramic bowl (8cm in diameter, 5cm in height) were used for object recognition; they were too heavy to be displaced by the animal and located at two corners of the apparatus. The mice did not show any preference for either of the two objects. For the implantation of cannulas, mice were first deeply anesthetized with ketamine (95mg/kg) and xylazine (10mg/kg) via intraperitoneal injections and stereotaxically implanted with a 26-gauge bilateral guide cannula (2mm long below pedestal, Plastic One, Roanoke, VA, USA) into the hippocampus using the following coordinates from Bregma: antero-posterior −2mm; medio-lateral +/-1.5mm; dorso-ventral −1mm from cortex. This was then held in place with dental cement. After surgery, animals were re-housed in home cage to recover at least for 7 days with close monitoring. Behavioral testing was conducted during the light phase of the 12h light/dark cycle under dim illumination (50lux). Prior to the first NOR test, during three consecutive days animals were handled 1min per day. On the day preceding each NOR test, animals were allowed to freely explore the empty (without objects) apparatus for 10min for habituation. The NOR test consisted of 2 trials (T1 and T2) separated by an inter-trial time interval (ITI). To assess the effect of acutely blocking Cx43 hemichannels and rescue by glutamine on NOR performance, mice were administered a bilateral intra-hippocampal injection (1μL per side; 1μL/min) of either Gap26 (1mM), Gap26 scramble (1mM) or Gap26 (1mM) + glutamine (200mM) 30min before T1 (n = 10 per group). This was carried out using a precision pump with a 33-gauge bilateral cannula (Plastics One) extending 0.5 mm below the end of the guide cannula to target hippocampus CA1 stratum radiatum (from Bregma: antero-posterior −2mm; medio-lateral +/-1.5mm; dorso-ventral −1.5mm from cortex). On T1 (acquisition trial), subjects were placed in the apparatus containing two identical objects (F1 and F2) for 10min before being returned to their home cage for an ITI of 24h. Then, they were placed back in the NOR apparatus containing a familiar (F) and a novel object (N) for 5min (T2, restitution trial). The type (familiar or novel) and the position (left or right) of the two objects was counterbalanced and randomized within each experimental group during T2. Between each trial, the NOR apparatus was cleaned with water and the objects with 70% ethanol. Exploration was defined as the animal directing the nose within 0.5cm of the object while looking at, sniffing or touching it, excluding accidental contacts with it (backing into, standing on the object, etc.). The raw exploration data were normalized to the total objects exploration time.

### Statistical analysis

Statistical analysis were performed using GraphPad Prism software (USA). All data are expressed as mean ± standard error of the mean (SEM), and n represents the number of independent experiments, unless otherwise stated. Statistical significance was determined by one-sampled t-test, two-tailed unpaired Student’s t-tests, one-way or two-way ANOVA with Bonferroni or Sidak’s post hoc tests. The test for normality was performed using Shapiro-Wik normality test. Individual data points are plotted in all bar graphs with p-values given in figure legends.

### Drugs

Gap26 (VCYDKSFPISHVR), and Gap26 scramble (PSFDSRHCIVKYV) were synthesized by Thermo Fisher Scientific (purity, 95%), and all others products were from Sigma unless otherwise stated.

### Data availability

We confirm that all relevant data are included in the paper and/or its supplementary information files.

## Acknowledgments

We thank P. Ezan, D. Mazaud and. J. Cazères for technical assistance. This work was supported by grants from the European Research Council (Consolidator grant #683154) and European Union’s Horizon 2020 research and innovation program (Marie Sklodowska-Curie Innovative Training Networks, grant #722053, EU-GliaPhD) to N.R. and from FP7-PEOPLE Marie Curie Intra-European Fellowship for career development (grant #622289) to G.C..

## Author Contributions

Conceptualization: N.R., G.C.; Data curation: G.C., N.R.; Formal analysis: G.C., J.M., G.D, O.C.; Funding acquisition: N.R., G.C.; Investigation: G.C., D.B., J.M., G.D., J.V., A.R., O.C.; Methodology: G.C., D.B., N.K., J.L.; Resources: N.K., C.M., A.B., I.L.; Supervision: N.R., I.L., J.L.; Project administration: N.R.; Validation: G.C., D.B., N.R.; Visualization: G.C., C.M., G.D., N.R.; Writing – original draft: G.C., N.R.; Writing – review & editing: G.C., N.R..

## Additional Information

Supplementary Information is available for this paper. Correspondence and requests for materials should be addressed to.

## Supplementary Information

**Supplementary Fig. 1.**
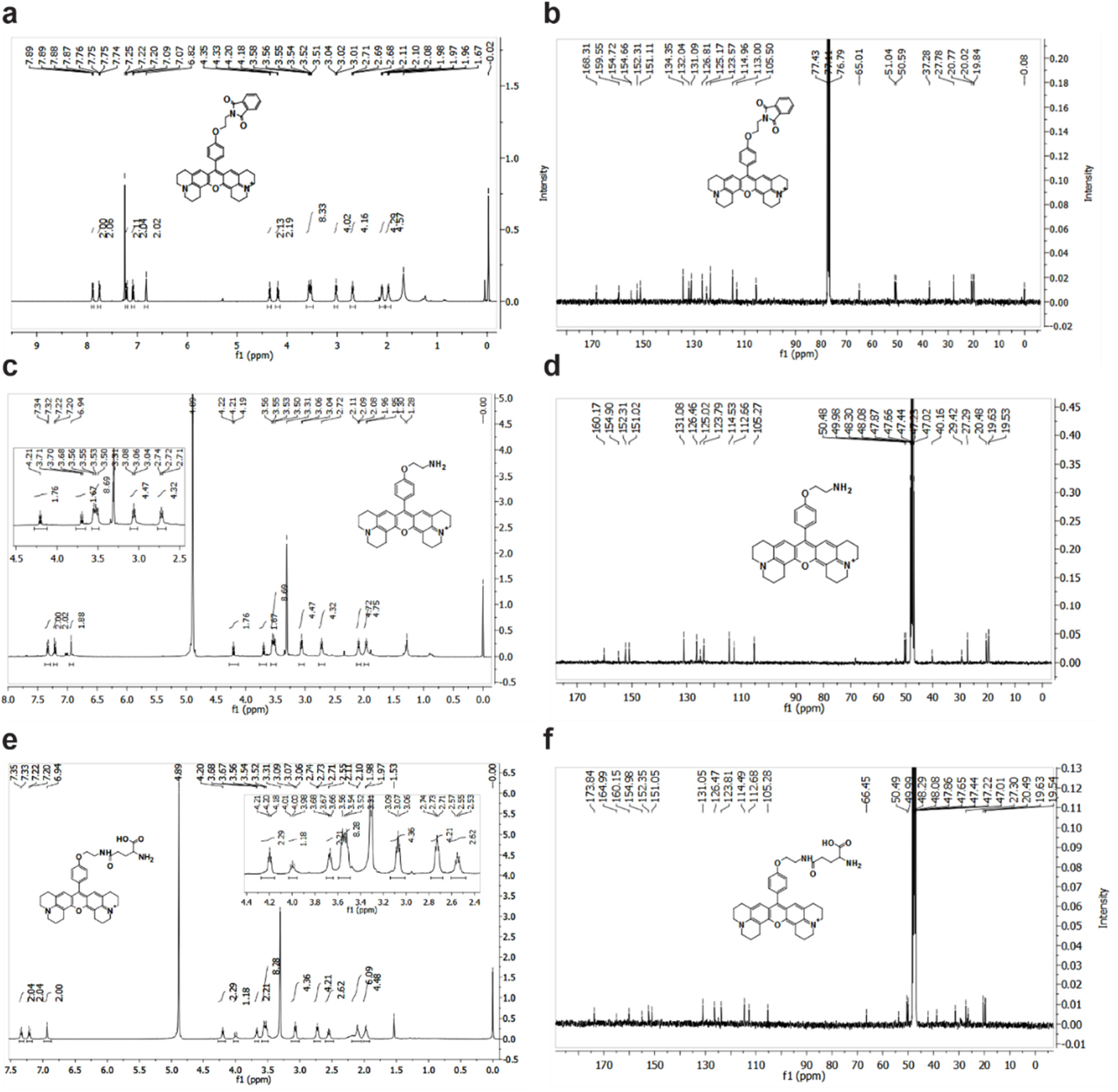
Structural confirmation of intermediate compounds 3, 4, and RhGln. **a,** ^1^H NMR (CDCl_3_, 400 MHz, 298 K and **b,** ^13^C NMR (CDCl_3_, 100 MHz, 298 K)) spectrum of compound **3** is shown. **c,** ^1^H NMR (CD3OD, 400 MHz, 298 K) spectrum of compound **4** is shown. Inset displays a zoom on the 4.5–2.5ppm region and **d,** ^13^C NMR (CD3OD, 100 MHz, 298 K) spectrum of compound **4**. **e,** ^1^H NMR (MeOD, 400MHz, 298 K) spectrum of RhGln is shown. Inset displays a zoom on the 4.4–2.4ppm region and **f,** ^13^C NMR (MeOD, 100 MHz, 298 K) spectrum of RhGln.

**Supplementary Fig. 2.**
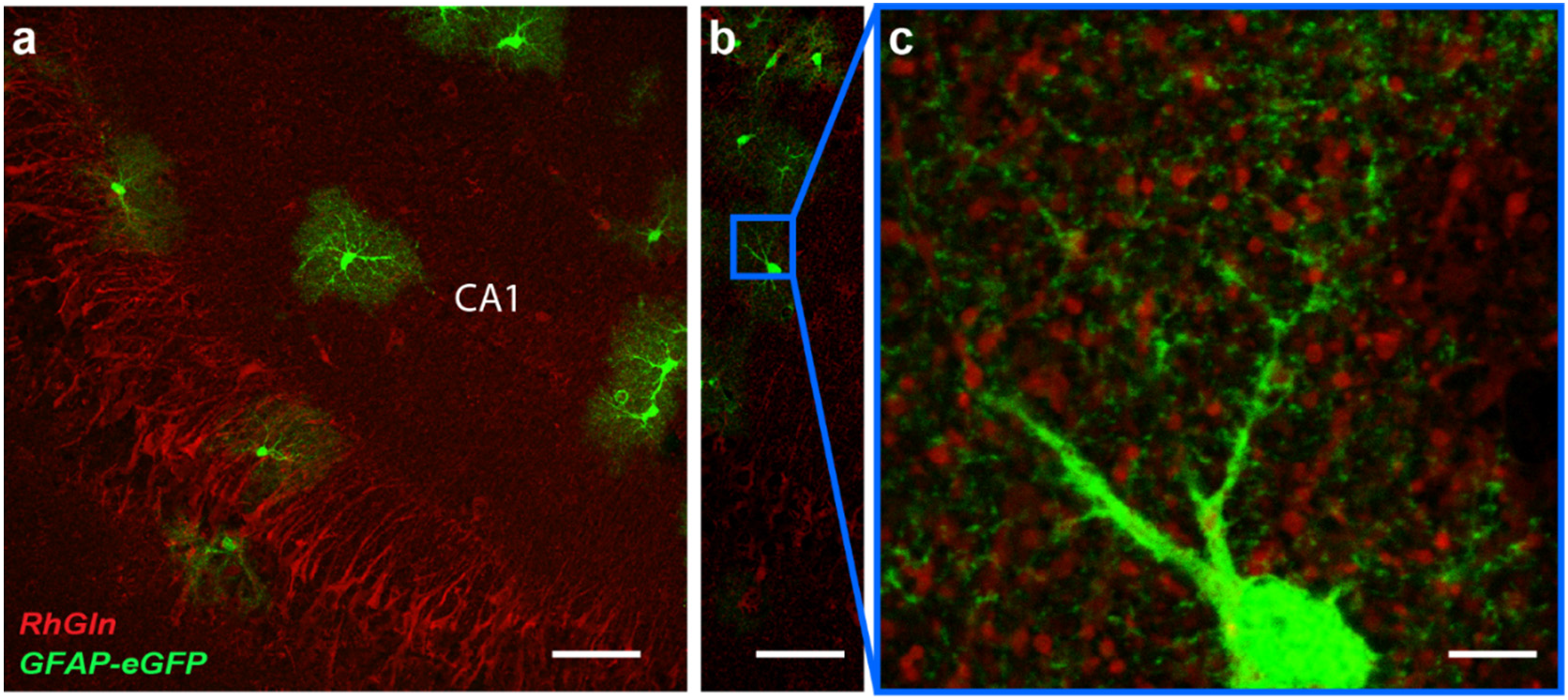
Extracellular bulk loaded RhGln does not enter astrocytes. Acute hippocampal slices obtained from GFAP-eGFP mice were loaded extracellularly with RhGln (0.05mM, 20min). Confocal images are shown in **(a-b)** low magnification of CA1 region and **(c)** high magnification of region marked by blue square in **b** containing astroglial processes (GFAP-eGFP positive, green), and RhGln (red). RhGln labeling does not co-localize with GFAP-eGFP expressing astrocytes. Scale bar: **a and b,** 50μm; **c,** 5μm.

**Supplementary Fig. 3.**
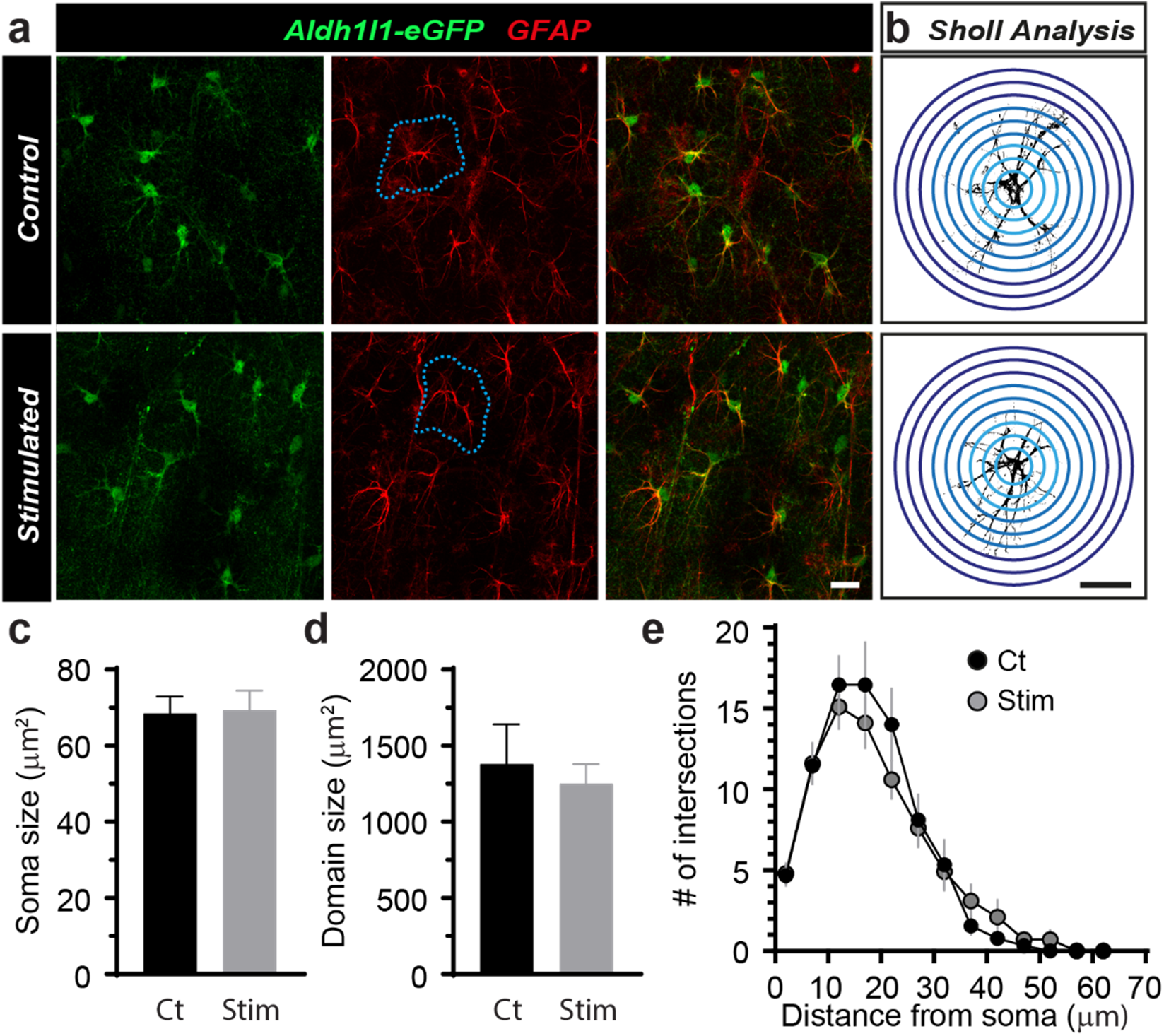
Morphology of astrocytes is unchanged upon stimulation of hippocampal slices. **a,** Representative images are shown for acute hippocampal slices obtained from Aldh1l1-eGFP mice with or without stimulation of the Schaffer collaterals (10Hz 30s every 3min for 20 min). Aldh1l1-eGFP expression is shown in green and immunostaining for GFAP is shown in red. **b,** Sholl analysis was performed as illustrated, where the number of intersections is measured from thresholded GFAP intensity over concentric circles drawn from the cell soma. **c-d**, Soma and domain sizes were measured and quantified from Aldh1l1-eGFP positive somas and GFAP immunostaining, respectively (n=60 cells from 3 independent experiments, *p*<0.999, Unpaired Student’s t-test). Domain size was determined by manually drawing an area encompassed by an entire astrocyte (blue dotted line in **a**). Quantification of Sholl analysis in **e** showed no significant stimulation-induced difference in astroglial morphology (Ct, n= 9; Stim, n=10, *p*=0.7129, Two-way ANOVA). Scale bar: **a and b**, 20μm.

